# Coverslip Hypoxia High-Content Screening (CH-HCS): A 2D Imaging Platform for Spatially Resolved Analysis of CAR-T Function Under Oxygen Gradients

**DOI:** 10.64898/2026.01.30.702833

**Authors:** João Pedro Zanon Murarolli, Giovana Beatriz Capuz Moreto, Maria Luiza Arrojo, Amanda Mizukami Martins, Sâmia Rigotto Caruso, Maristela Delgado Orellana, Rodrigo Alexandre Panepucci

## Abstract

Hypoxia within the tumor microenvironment profoundly limits the efficacy of immune and cellular therapies, yet most in vitro cytotoxicity assays neglect spatial oxygen heterogeneity. We developed Coverslip Hypoxia High-Content Screening (CH-HCS), a simple, scalable 2D co-culture platform that enables quantitative, region-resolved evaluation of CAR-T cell activity across controlled oxygen gradients within a single well. In CH-HCS, a 5 mm glass coverslip placed over tumor-stroma co-cultures restricts oxygen diffusion, generating concentric hypoxia–normoxia zones in standard 96-well plates. Fluorescently labeled CD19^+^ Raji or CD19^−^ K562 tumor cells, stromal cells, and anti-CD19 CAR-T cells were analyzed using multiparametric fluorescence imaging with an ImageXpress Micro XLS system coupled to custom CellProfiler–KNIME pipelines, enabling segmentation of spatial Regions of Interest—InnerCore, OuterCore, Periphery, and Outside—and single-cell quantification of tumor death (SYTOX Green) and T-cell morphodynamics. The platform reproducibly established oxygen gradients that strongly shaped cellular behavior: CAR-T cytotoxicity and motility were maximal in normoxic regions but markedly suppressed within hypoxic cores, whereas effector cell survival increased under low oxygen. Unlike bulk cytotoxicity assays, CH-HCS directly visualizes spatial functional heterogeneity within the same well, allowing simultaneous comparison of matched hypoxic and normoxic compartments. Together, CH-HCS provides a cost-effective, high-throughput, and physiologically relevant tool for preclinical screening of CAR-T products and therapeutic strategies aimed at overcoming hypoxia-driven immune resistance at the tumor–stroma interface.

**Key Point:**

- **Spatially resolved quantification:** Simultaneous measurement of cellular behavior in matched hypoxic and normoxic compartments within the same well.
- **Physiologically relevant gradients:** Coverslip geometry generates reproducible oxygen diffusion profiles that emulate the in vivo tumor–stroma interface.
- **Multiparametric single-cell readouts:** High-content imaging coupled with automated segmentation provides data on viability, cytotoxicity, and effector morphology.
- **High-throughput and low-cost:** Fully compatible with 96-well formats, enabling parallel pharmacological or genetic screening without specialized hypoxia chambers.
- **Generalizable design:** Applicable beyond CAR-T assays to study stromal adaptation, drug resistance, or immune suppression across controlled oxygen gradients.

## 1 Introduction

The discovery that T-cell activity is regulated by immune checkpoint molecules such as PD-1, PD-L1, and CTLA-4^(*1*)^, and that these pathways can be co-opted by cancers as immune evasion mechanisms within the tumor microenvironment (TME)^(*2-4*)^, led to the development of Checkpoint Inhibitor Therapies (CITs), which have revolutionized cancer treatment. CITs are based on monoclonal antibodies (MoAbs) that target and block these inhibitory checkpoint molecules, thereby promoting strong anti-tumor immune responses^(*5-7*)^. Adoptive Cell Therapies (ACTs) have also revolutionized cancer treatment^(*8-10*)^. The development of engineered Chimeric Antigen Receptors (CARs), combining TCR structural and signaling components with a variable antigen-binding domain derived from a monoclonal antibody targeting the CD19 B-cell marker, enabled genetically modified anti-CD19 CAR-T cells to elicit robust immune responses against relapsed and refractory B-cell malignancies, including Non-Hodgkin lymphomas (NHL)^(*9, 11-18*)^.

Despite their revolutionary role in cancer treatment, the efficacy of CITs and ACTs depends on the ability of immune cells to overcome distinct barriers within the TME. For instance, a pre-existing intratumoral adaptive immune response is crucial for the effectiveness of CITs^(*19*)^. Similarly, in the case of ACTs, infused CAR-T cells must infiltrate the tumor to engage in direct cytotoxic killing of target cancer cells, a requirement that largely limits their efficacy in solid tumors^(*20-22*)^. Tumors with a pre-existing immune response are referred to as Inflamed or Hot tumors, whereas those poorly infiltrated by lymphocytes are referred to as Non-Inflamed or Cold tumors^(*23*)^. Among the latter, Immune-Silent tumors exhibit a near-complete absence of immune cells, likely reflecting defective chemoattraction, whereas in Immune-Excluded tumors immune cells are restricted to the periphery of tumor nests by local mechanical and/or functional barriers that prevent infiltration and suppress immune cell function and survival^(*22,24, 25*)^. Infiltration by NK cells, cytotoxic CD8+ T cells, helper CD4+ Th1 lymphocytes, and M1 tumor-associated macrophages (TAMs) is associated with favorable prognosis, whereas the presence of regulatory T cells (Tregs) and M2 TAMs correlates with poor clinical outcomes^(*19*)^. Likewise, immune cell presence within affected lymph nodes (LNs) prior to therapy, as well as the accumulation of infused anti-CD19 CAR-T cells, influences treatment response in B-NHLs^(*26-31*)^. Thus, the same TME barriers that restrict the infiltration and function of endogenous immune cells also represent a major obstacle to immunotherapies in general, limiting the infiltration, trafficking, and function of both endogenous and adoptively transferred immune cells, including CAR-T cells^(*32-36*)^.

Hypoxia-driven metabolic reprogramming of cells in the TME can affect recruitment, infiltration, survival, activation and function of cancer, stromal and immune cells, thus directly impacting the immune landscape and anti-tumor response^(*37-41*)^. For example, hypoxia promotes M2 polarization of myeloid and stromal cells and the generation of Tregs^(*42-45*)^, induces a shift from Th1 to Th2 responses^(*46*)^, and drives the expression of immune checkpoint molecules in T^(*47*)^ and NK cells^(*48, 49*)^. Nutrient deprivation and lactate accumulation also dramatically hinders T^(*42, 50*)^ and NK cell function^(*51*)^, while conferring a selective advantage to Tregs^(*52, 53*)^. Hypoxia-inducible factors (HIFs) are master regulators of tumor progression, inducing the transcription of genes that promote angiogenesis, extracellular matrix remodeling, epithelial-mesenchymal transition, motility, invasion, metastasis, cancer/leukemic stem cells (C/LSCs) maintenance, immune evasion, resistance to chemo- and radio-therapy^(*54*)^, as well as metabolic reprogramming^(*55, 56*)^. Tumor-associated stromal cells, including mesenchymal stromal cells (MSCs) and cancer-associated fibroblasts (CAFs), are major components of the TME niche, promoting tumor progression while constituting a barrier to immune cell entry and activity^(*41, 57-62*)^. The hypoxic-stromal niche of the bone marrow^(*63, 64*)^, the spleen and LNs, also promote stemness of normal and C/LSCs, constituting classic relapse sites^(*65-72*)^.

Adenosine signaling downstream of hypoxia mediates several immune evasion mechanisms in the TME ^(*73, 74*)^. Hypoxia-activated HIF-1α induces the transcription of the enzymes CD39 and CD73 (a MSC marker), which converts extracellularly released ATP into Adenosine^(*75-78*)^. Hypoxia also downregulates the expression of Equilibrative Nucleoside Transporters (ENTs) involved in the cellular uptake of adenosine, further promoting its accumulation^(*74, 79, 80*)^. In turn, adenosine is rapidly converted into inosine by Adenosine DeAminase (ADA)^(*81*)^, which can be anchored in the membrane by CD26, protecting cells from adenosine’s functions^(*82*)^. Adenosine signaling via its A2A receptor strongly suppresses CD8+^(*83*)^, Th1 and Th17 effector T-cells, and induce Tregs generation^(*84*)^. Also, A2B signaling promotes metastatic behavior of tumor cells^(*85-87*)^. Overall, hypoxia-adenosinergic signaling is an attractive target in the TME^(*20, 88, 89*)^ and several reports have showed that the resistance to chemotherapy and immunotherapy of CD39^High^ and CD73^High^ tumors can be bypassed with the combined use of antagonists of A2A, CD39, or CD73, driving a series of pre-clinical and clinical trials^(*90-94*)^. Understanding how hypoxia-adenosinergic signaling impacts the distribution of cancer, stromal, and immune cells within the TME may provide important insights for the development of new therapeutic strategies^(*95*)^.

Despite ongoing efforts to develop physiologically relevant “immunocompetent” in vitro systems that model cellular heterogeneity and metabolic and oxygen gradients within the TME^(*96-98*)^, suitable and straightforward experimental approaches remain scarce. In this context, modern imaging techniques have shown an enormous potential as tools to dissect the TME^(*99, 100*)^, in 2D monolayer^(*101-104*)^ and 3D spheroid models^(*105-111*)^.

In this study, we present a 2D tumor–stroma model that imposes controllable hypoxia gradients and couples them with automated, quantitative fluorescence imaging to evaluate CAR-T function. Our Coverslip Hypoxia High-Content Screening (CH-HCS) approach resolves spatial compartments within each well, enabling real-time, region-specific quantification of tumor (e.g., Raji) and effector (CAR-T) cell proliferation, death, and activation under matched hypoxic vs. normoxic conditions—serving as an in-plane analogue of a spheroid cross-section—and providing a practical, high-throughput platform to interrogate TME-imposed barriers and test strategies to overcome them.

## 2 Material and Methods

### 2.1 Cell Lines, Primary Cells, and Culture Conditions

Human Umbilical Cord Multipotent Mesenchymal Stem/Stromal cells (UC-MSCs) were cultured in α-MEM supplemented with 10 % fetal bovine serum (FBS) and 1 % penicillin/streptomycin at 37 °C and 5 % CO_2_. For co-culture assays, UC-MSCs cells were seeded at 1 × 10^4^ cells per well (in a total volume of 100µl of the α-MEM medium) in 96-well black, clear-bottom plates (Cost ar 3603, Corning) and allowed to adhere for 24 h, forming a near-confluent stromal monolayer (Fig. 1, step 1).

**Figure 1.**
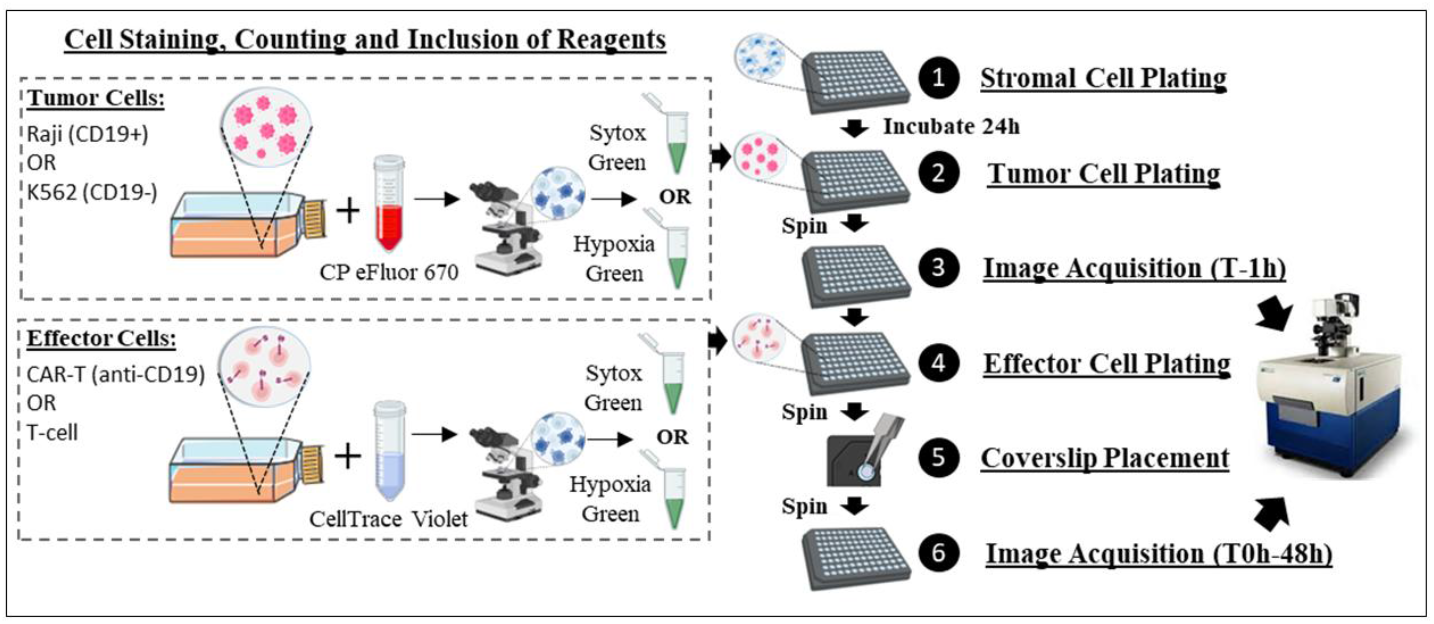
Stepwise workflow of the Coverslip Hypoxia High-Content Screening (CH-HCS) assay. (1) UC-MSCs are seeded in 96-well black, clear-bottom plates and cultured for 24 h to form a confluent stromal monolayer. (2) Tumor cells (Raji CD19^+^ or K562 CD19^−^) are harvested, stained with Cell Proliferation Dye eFluor 670 (CP670), resuspended in medium containing the viability dye SYTOX™ Green and/or the hypoxia reporter Image-iT™ Green, and plated on top of the stromal monolayer. (3) Following a brief plate centrifugation, baseline images are acquired (T−1 h). (4) Effector lymphocytes (anti-CD19 CAR-T or CAR-Jurkat) are harvested, stained with CellTrace™ Violet (CTV), resuspended in medium containing SYTOX™ Green or Image-iT™ Green, and added to wells already containing the tumor–stroma co-culture. (5) After another brief centrifugation, a 5 mm glass coverslip is gently placed at the center of each well and the plate is centrifuged again to ensure contact and uniform oxygen restriction. (6) Plates are imaged at sequential time points using an ImageXpress Micro XLS high-content microscope. The acquired images are processed through integrated CellProfiler–KNIME pipelines for spatial segmentation (Inner Core, Outer Core, Periphery, Outside) and single-cell quantification of viability, hypoxia, and effector morphology.

Raji cells (Burkitt lymphoma, CD19^+^; ATCC CCL-86) ^(*112-114*)^, K562 cells (myelogenous leukemia, CD19^−^), and Jurkat E6-1 cells (ATCC TIB-152), were maintained in RPMI-1640 (Thermo Fisher) containing 10 % FBS, 1 % penicillin/streptomycin, and 25 mM HEPES.

### 2.2 CAR-Jurkat and Jurkat Control Cells

To validate the 2D heterotypic CH-HCS system, anti-CD19 CAR-Jurkat cells were used as a standardized model. This system enabled the assessment of hypoxia, cytotoxic function, activation, and survival against CD19^+^ targets (Raji) or control CD19^−^ K562 cells. Non-transduced Jurkat E6-1 cells served as negative controls. The anti-CD19 CAR-Jurkat cell line was derived from Jurkat E6-1 cells (ATCC TIB-152) ^(*115, 116*)^ transduced with the pCAR19 lentiviral vector, encoding a chimeric antigen receptor (CAR) containing an anti-CD19 scFv and a 4-1BB co-stimulatory domain.

Following transduction, fluorescence-activated cell sorting (FACS) was performed to isolate cells expressing the CAR construct. Surface expression of the CAR was subsequently confirmed by flow cytometry (Annex I) using Alexa Fluor 647 AffiniPure F(ab’)_2_ Fragment Goat Anti-Mouse IgG, F(ab’)_2_ fragment-specific (Jackson Immuno Cat. 115-606-072) and its isotype control Alexa Fluor 647 ChromPure Goat IgG F(ab’)_2_ Fragment (Cat. 005-600-006). Data were analyzed using FlowJo v10.

### 2.3 Primary CAR-T Cells and T-Cell Controls

After establishing the CH-HCS system, primary anti-CD19 CAR-T cells were subsequently introduced into the model to enable functional assessment. These CAR-T cells were generated by lentiviral transduction of immunomagnetically isolated human T cells using a second-generation CAR vector containing CD3ζ/4-1BB coestimulatory domains (ProBio - GenScript Biotech Corporation). Peripheral T lymphocytes were isolated from healthy-donor leukapheresis samples under informed consent (TCLE) and used as the source material for CAR-T generation under validated research protocols. Within each experiment, CAR-T cells and non-transduced T-cell controls were derived from the same donor, ensuring donor-matched conditions. The culture was maintained in RPMI supplemented with 10% fetal bovine serum (FBS), 1 % penicillin/streptomycin and 100 U/mL IL-2 at 37 °C and 5 % CO_2_.

Following incorporation into the CH-HCS model, CAR-T cells and non-transduced T cells (negative controls) were employed to evaluate spatial behavior, activation, cytotoxicity, and hypoxia-dependent viability. CAR expression was confirmed by flow cytometry (Annex II) using the same detection reagents as above, and data were analyzed in FlowJo v10, confirming consistent CAR expression prior to functional assays.

### 2.4 Cell Labeling and Preparation for the Coverslip-Hypoxia Assay

To enable cell-type discrimination, Tumor cells (Raji and K562) were stained with Cell Proliferation Dye eFluor 670 (Invitrogen 65-0840-85; Ex/Em 630/670 nm) at 4 µM, whereas Effector T cells (CAR-Jurkat/CAR-T and control Jurkat and T cells) were stained with CellTrace™ Violet (CTV) (Invitrogen C34557; Ex/Em 405/450 nm) at 1 µM. After staining, cells were incubated in complete RPMI supplemented with 10 % FBS for 3-4 h (37 °C, 5 % CO_2_) to allow cell recuperation.

Tumor or Effector T cells were independently resuspended at densities of 4×10^5^ cells/mL or 8×10^5^ cells/mL, respectively, in RPMI medium containing SYTOX™ Green (Invitrogen S7020;, Ex/Em 504/523 nm) at a 40 nM concentration. After removing 50µl of the α-MEM medium from each well containing the UC-MSC monolayer, 25µl of the Tumor cell suspension were dispensed per well (1×10^4^ cells per well) over the UC-MSCs monolayer (Fig. 1, step 2). Plates were centrifuged at 1,200 rpm (≈150 ×g) for 3 min and imaged for baseline acquisition (T −1 h) (Fig. 1, step 3). Subsequently, 25µl of the Effector T cell suspension were dispensed per well (2×10^4^ cells per well), with a final concentration of SYTOX Green of a 20 nM (Fig. 1, step 4). Plates were centrifuged again under identical conditions and, immediately after, a 5 mm glass coverslip was gently placed in each well, using fine point tweezers (Fig. 1, step 5), and the plate was centrifuged again, prior to the T0 acquisition (Fig. 1, step 6).

In selected experiments, tumor and effector T cells were resuspended in medium containing the Image-iT™ Green Hypoxia Reagent (Invitrogen I14833; Ex/Em 488/520 nm) at 2 µM and plated in parallel wells (final concentration of 1 µM) to enable functional validation of hypoxic conditions. The Image-iT signal was used to evaluate the percentage of hypoxic cells within the intrawell microenvironment.

To evaluate the role of adenosinergic signaling, wells were treated with the A_2_A receptor antagonist Ciforadenant /CPI-444 (MedChem HY-101978) ^(*117*)^ or 0.1 % DMSO vehicle. In this experiment, α-MEM was completely removed from the wells with the stromal monolayer and replaced with 50µl of α-MEM containing Ciforadenant at 80nM. After 4h, Tumor and Effector cells were added, attaining a final concentration of 40nM, and following coverslip placement, images were acquired (T_0_). CAR-T cytotoxicity was quantified across all oxygen-defined regions after 40 h (Fig. 6).

### 2.5 Coverslip Hypoxia (CSH) assay

To impose controlled radial oxygen gradients in 2D monolayers, we implemented the Coverslip Hypoxia (CSH) system (Fig. 1). Fluorescently labeled tumor cells were plated on pre-seeded UC-MSCs monolayers in black, clear-bottom 96-well plates (Corning Costar 3603, with a well bottom diameter of 6.35mm). After gentle centrifugation (150 × g, 5 min) to promote contact, a sterile 5 mm circular glass coverslip was placed at the center of each well, physically restricting O_2_ diffusion and creating hypoxic (beneath the coverslip) and normoxic (outside) compartments. Cell densities were optimized to achieve a uniform stromal cell monolayer at the well bottom and a reproducible hypoxia gradient beneath the coverslip, while keeping an adequate separation of tumor and effector cells, to allow a clear segmentation.

Real-time reporters were added immediately before incubation (in separate wells): Image-iT™ Green Hypoxia reagent (1.25 µM; Invitrogen I14834) for hypoxia mapping, and SYTOX™ Green (1 µM; Invitrogen S7020) to detect membrane-compromised cells. Plates were imaged at 0 and 24h (with additional imaging at 12 and 40 h for hypoxia and drug-response experiments) using three technical replicates per condition (three wells per experimental condition). The configuration produced four spatial oxygenation zones: InnerCore, OuterCore, Periphery, and OutsideCoverslip. The CSH setup, region nomenclature, and radial diffusion concept are schematized in Fig. 2.

**Figure 2.**
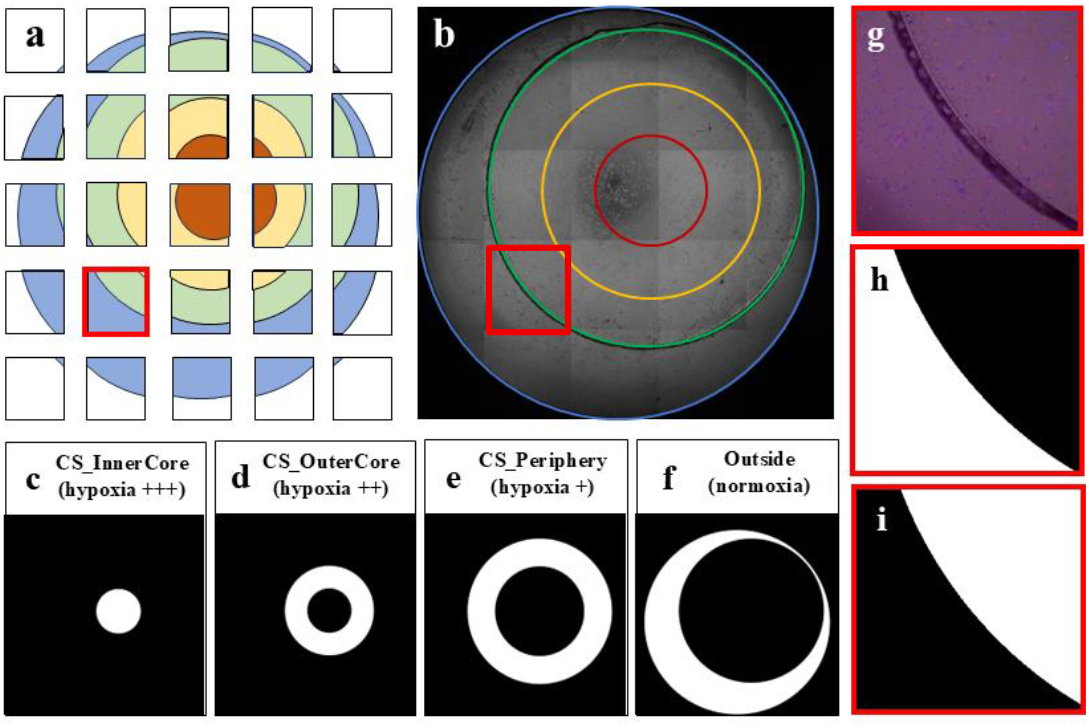
Automated processing and segmentation of Regions of Interest (ROIs) in CellProfiler for the quantitative analysis of hypoxic and normoxic zones in CSH model. The CellProfiler pipeline processed tiled fluorescence images from 96-well plates to define spatial zones corresponding to distinct oxygenation levels beneath and outside the coverslip. (a) Illustrative schematic of the acquisition mosaic showing the defined Regions of Interest (ROIs): *OutsideCoverslip* (blue), *Periphery* (green), *OuterCore* (yellow), and *InnerCore* (red). (b) Composite transmitted-light (TL) image with the same spatial segmentation displayed in (a). (c–f) Sequential segmentation steps using CellProfiler-generated masks: (c) *InnerCore* (severe hypoxia), (d) *OuterCore* (moderate hypoxia), (e) *Periphery* (transition zone), and (f) *Outside* (normoxic region). The red rectangle highlights an individual field located at the coverslip interface. (g) Cropped object restored to its original scale, overlaid with TL, DAPI (CAR-T), and Cy5 (Raji) channels, enabling assignment of segmented cells to their corresponding spatial compartment: (h) *Outside* or (i) *Periphery*.

### 2.6 High-Content Multiparametric Fluorescence Microscopy (HCS)

Quantitative analysis was performed using a High-Content Screening (HCS) approach combining automated fluorescence microscopy and computational image quantification. Images were acquired using an ImageXpress MICRO XLS system (Molecular Devices) equipped with a transmitted light (TL) tower and five excitation/emission filter sets: DAPI-5060C (Ex 377/50; Em 447/60; blue); FITC-3540C (Ex 482/35; Em 536/40; green); Cy3-4040C (Ex 531/40; Em 593/40; orange); TxRed-4040C (Ex 562/40; Em 624/40; red), and Cy5-4040C (Ex 628/40; Em 692/40; deep red). The DAPI channel was used to image CTV-stained Effector T cells, FITC channel to detect Image-iT hypoxia and/or SYTOX death, and the Cy5 channel to acquire images of CP670-labeled tumor cells.

After establishment of the optimal focal plane and exposure time for each fluorescence channel, image acquisition was performed using MetaXpress software version 5.3 (Molecular Devices) with these settings maintained consistently across all subsequent time points. All acquired images were exported as independent TIFF files for each channel (TL, DAPI, FITC, and Cy5). Acquisitions executed at different time points were exported using distinct Plate Names. Each well was imaged as a 5 × 5 field grid (25 fields per well) at 10× magnification (field of view ≈ 1.4 × 1.4 mm), ensuring complete coverage of the well area and adequate resolution to resolve spatial oxygen-gradient zones (Figs 2–3). During image acquisition, plates were maintained under controlled temperature conditions, with no active CO_2_ control. Representative acquisitions and overlays used for segmentation are shown in Fig. 2.

### 2.7 Automated Image Analysis

#### 2.7.1 General Workflow

An overview of the spatial segmentation logic and the final association of single-cell objects with their corresponding oxygen-defined regions is illustrated in Fig. 2– 4.

Quantitative image processing and spatial mapping of the Coverslip Hypoxia High-Content Screening (CH-HCS) assay were performed using CellProfiler 4.2.8 through a structured workflow composed of five sequential pipelines (Supplementary Fig. S1). Results produced by each pipeline were saved in dedicated output folder directories and metadata tables, enabling full traceability and reproducibility from raw image acquisition to per-cell quantitative datasets.

The complete workflow consisted of the following pipelines (and output folder):

1. **Pipeline_01 – Image Tiling and Metadata Extraction (Output_01_TiledImages);**
2. **Pipeline_02 – Segmentation of Well and ROI Objects (Output_02_Objects)**
3. **Pipeline_02b – Plate-level quality control (Output_02b_TiledPlates)**
4. **Pipeline_03 – ROI cropping and registration (Output_03_CroppedObjects); and**
5. **Pipeline_04 – Single-cell segmentation and feature quantification (Output_04_Results)**.

All images acquired by the ImageXpress MICRO XLS system were exported as independent TIFF files for each channel (TL, DAPI, FITC, and Cy5). The five pipelines were executed sequentially as shown in Supplementary Fig. S1.

#### 2.7.2 Pipeline 01 – Image Tiling and Metadata Extraction

The first pipeline (*Pipeline_01_Tile-WellImages_Images-TiledImages.cpproj*) performed preprocessing and reconstruction of full-well composite images with the TL images, to allow segmentation of the well and coverslip-associated regions (Supplementary Fig. S2).

To ensure proper sequential assembly of composite images in CellProfiler, a PowerShell script was implemented to automate the renaming of image site identifiers across multiple experiment folders exported by the MetaXpress acquisition software (Supplementary Script 1). The script iteratively scanned subdirectories containing multi-site image sets and reassigned standardized site identifiers within filenames (e.g., *s1* → *s01*) to maintain compatibility with CellProfiler’s metadata parsing system. Key modules included:

i. **Metadata** – regular expressions captured Plate Name, Well, Site, and Channel n° from filenames.
ii. **Resize** – reduced image resolution to 12.5 % of the original to optimize computation.
iii. **Tile** – assembled the 5×5 (25 site) image grid into a whole-well composite image.
iv. **NamesAndTypes** and **Groups** – defined channel name and grouped images by plate and well.
v. **SaveImages** – exported stitched TIFFs to the directory *Output_01_TiledImages*.

These composite images preserved the spatial relationship between the coverslip, well edges, and all acquired fields, providing the structural basis for ROI segmentation.

#### 2.7.3 Pipeline 02a – Plate-Level Quality Control

Before proceeding to segmentation of the well regions, plate-level quality control was performed using *Pipeline_02b_TilePlates_QC.cpproj* (Supplementary Fig. S3). This pipeline generated composite images of all acquired wells within a plate, enabling direct comparison of stitched bright-field images captured at the beginning (0 h) and end (48 h) of the experiment. The procedure verified coverslip positioning and excluded potential drift arising from incubation or repeated imaging. Once positional stability was confirmed, only a single time point (typically 0 h) was used for final ROI segmentation.

#### 2.7.4 Pipeline 02b – Segmentation of Well and ROI Objects

The second pipeline (*Pipeline_02_Segment-WellObjects_TiledImages-Objects.cpproj*) generated spatial regions of interest (ROIs) corresponding to the distinct oxygenation zones beneath and outside the coverslip (Supplementary Fig. S4). The sequence of ROI generation is illustrated in **Supplementary Fig. S5** and in Figure 2. Processing steps included:

i. **Well Object –** The entire illuminated region of the well was automatically segmented using the *IdentifyPrimaryObjects* module, which uses the transmitted-light image to detect the circular well border, exploiting the contrast of black-walled plates. This region represents the total analysis area for the subsequent Boolean operations
ii. **Coverslip (CS) Object –** The hypoxia-inducing coverslip was manually defined for each well using the *Crop* module with an elliptical selection tool. The cropping window was centered on the coverslip’s position and saved as *CoverslipCrop*. This region represents the physical barrier that limits oxygen diffusion and establishes the hypoxic gradient.
iii. **CS_InnerCore ROI** – To generate the most hypoxic compartment, the cropped coverslip image (CoverslipCrop) was reduced to its central pixel using *ShrinkToObjectCenters* module, creating the CoverslipCenter, which was them expanded by 150 pixels by the *ExpandOrShrinkObjects* module to define the InnerCore, corresponding to severe hypoxia at the center of the coverslip.
iv. **CS_OuterCore ROI** – The InnerCore object was expanded by 150 pixels using the *ExpandOrShrinkObjects* module to generate the Core region. The OuterCore was then defined by subtracting the InnerCore from the Core region using the *IdentifyTertiaryObjects* module. These annular regions represent zones of moderate hypoxia surrounding the central core.
v. **CS_Periphery ROI** – The Periphery (transition zone) was obtained by subtracting the Core from the Coverslip object using the *IdentifyTertiaryObjects* module. This region corresponds to the hypoxia-to-normoxia interface where oxygen diffusion gradients decline sharply.
vi. **OutsideCoverslip ROI** – Finally, the OutsideCoverslip (normoxic region) was computed by subtracting the Coverslip area from the total Well object. This region served as the internal normoxic control for each well.

Through these hierarchical morphological and Boolean operations, four concentric ROIs— **InnerCore, OuterCore, Periphery**, and **OutsideCoverslip**—were consistently defined across all wells. This automated segmentation framework provided spatially standardized oxygen-gradient zones, forming the computational foundation for single-cell quantification and region-resolved analyses in the CH-HCS assay (see Fig. 2). The *MeasureObjectSizeShape* module was used to precisely determine the area corresponding to each ROI in every well, and the results were exported to a spreadsheet using the *ExportToSpreadsheet* module. Each segmented object was also saved as an 8-bit TIFF mask using the *SaveCroppedObjects* module, generating filenames that preserved the original tiled well transmitted-light (TL) image name followed by a suffix indicating the corresponding object. All output files—including numerical data and mask images—were stored in the directory *Output_02_Objects*.

#### 2.7.5 Pipeline 03 – ROI cropping and registration

The third pipeline (*Pipeline_03_Crop-WellObjects_Objects-CroppedObjects.cpproj*) subdivided the full-well composites into 25 rectangular sub-images that corresponded exactly to the original acquisition fields (Supplementary Fig. S6).

i. **Metadata / NamesAndTypes** modules imported the ROI masks, with the regular expression modified to capture not only the well identifier but also the object name appended as a suffix in the previous pipeline.
ii. The **Groups** module organized wells for batch processing.
iii. The **Crop** module divided each stitched object image according to predefined spatial coordinates, generating cropped ROI image files. Each cropped image received a suffix identifier corresponding to its site number, matching the original 5×5 tiled transmitted-light (TL) image.
iv. **SaveImages** exported the cropped TIFFs to *Output_03_CroppedObjects*.

These sub-images were later resized back to their native resolution and overlaid with fluorescence channels for single-cell segmentation. The procedure ensured that each cell could be linked to its precise spatial ROI and acquisition site (see red inset in Supplementary Fig. S6).

#### 2.7.6 Pipeline 04 – Single-cell segmentation and feature quantification

The fourth pipeline (*Pipeline_04_Analyse_CroppedObjectsAndImages-Results.cpproj*) constituted the final and most computationally intensive stage of the CH-HCS image-analysis workflow. It integrated all cropped object sub-images (25 sites per well) with the corresponding full resolution images from fluorescence—**DAPI, FITC, Cy5**— and **transmitted light** channels to perform single-cell segmentation, feature extraction, and spatial mapping of tumor and effector populations across the oxygen-defined regions of the Coverslip Hypoxia (CSH) model (Supplementary Figs. S7a–S7b). The workflow comprised:

i. **Data Loading and Metadata parsing** – The pipeline began by loading all fluorescence channels at full resolution together with the cropped ROI objects produced in *Pipeline 03* (imported as binary masks, with their original labels). In the **Metadata** module, regular expressions were used to extract *Plate, Well*, and *Site* identifiers from each image filename, while the cropped object filenames also provided the *Object* identifier corresponding to the specific ROI (InnerCore, OuterCore, Periphery, or Outside), as well as the corresponding site. These metadata entries were automatically cross-referenced with an experimental CSV metadata file (*MetaData.csv*) containing condition-level variables (well, cocultured cell types, drug, dose, replicate number, etc), ensuring that every image and segmented object was linked to its biological and experimental context. The metadata was later used to automate grouping and tracking across downstream image and data analysis stages carried in KNIME.
ii. **NamesAndTypes alignment** – In this module, channel identities were defined for each fluorescence image (DAPI, FITC, Cy5, TL), while cropped ROI masks were renamed with a sufix “_NotResized” (to distinguish from the later resized object). Images and object masks were aligned based on their corresponding identifiers (i.e. plate. well, and site, for image channels; and well and site, for the object masks, derived from a single timepoint).
iii. **RunCellPose modules** – GPU-accelerated segmentation of:
  a. *Effector T cells* (CTV^+^, DAPI channel),
  b. *T*umor cells (CP-670^+^, Cy5 channel), and
  c. *Nuclei from Dead cells* (SYTOX Green^+^, FITC channel) (**Fig. 4a–d**).
iv. **RelateObjects** and **ClassifyObject** – association of SYTOX^+^ nuclei (or Hypoxia Green+ cells) with parent Tumor or T-cell objects and classification as *live* or *dead (or Hypoxic or normoxic*.
v. **RelateObjects (ROI linkage)** – mapping of each cell to its spatial region (InnerCore, OuterCore, Periphery, Outside).
vi. **MeasureObjectIntensity / MeasureObjectSizeShape** – extraction of fluorescence intensities, morphological parameters (area, perimeter, eccentricity), and spatial coordinates.
vii. **ExportToDatabase** – consolidation of all cellular features and metadata into SQLite databases (*Output_04_Results*).

The resulting dataset contained per-cell measurements of viability, hypoxia, and morphology, all indexed by oxygen-region and acquisition site. This enabled spatially resolved quantification of tumor and effector populations across diffusion-defined compartments of the CH-HCS assay.

### 2.8 Single-cell segmentation, measurements, and derived readouts

Death was quantified by SYTOX positivity (FITC), using fixed intensity thresholds applied consistently across all conditions. Hypoxia was quantified by Image-iT intensity (FITC). Morphometric features (e.g., eccentricity) were computed per effector cell as a proxy for activation/motility states. All single-cell objects were related to their parent ROI (InnerCore, OuterCore, Periphery, OutsideCoverslip) to enable spatially resolved counts, percentages, and densities (cells/mm^2^). Representative spatial hypoxia maps and population trends are in Fig. 3; cytotoxicity (tumor death) and effector survival/activation are in Fig. 5–6.

**Figure 3.**
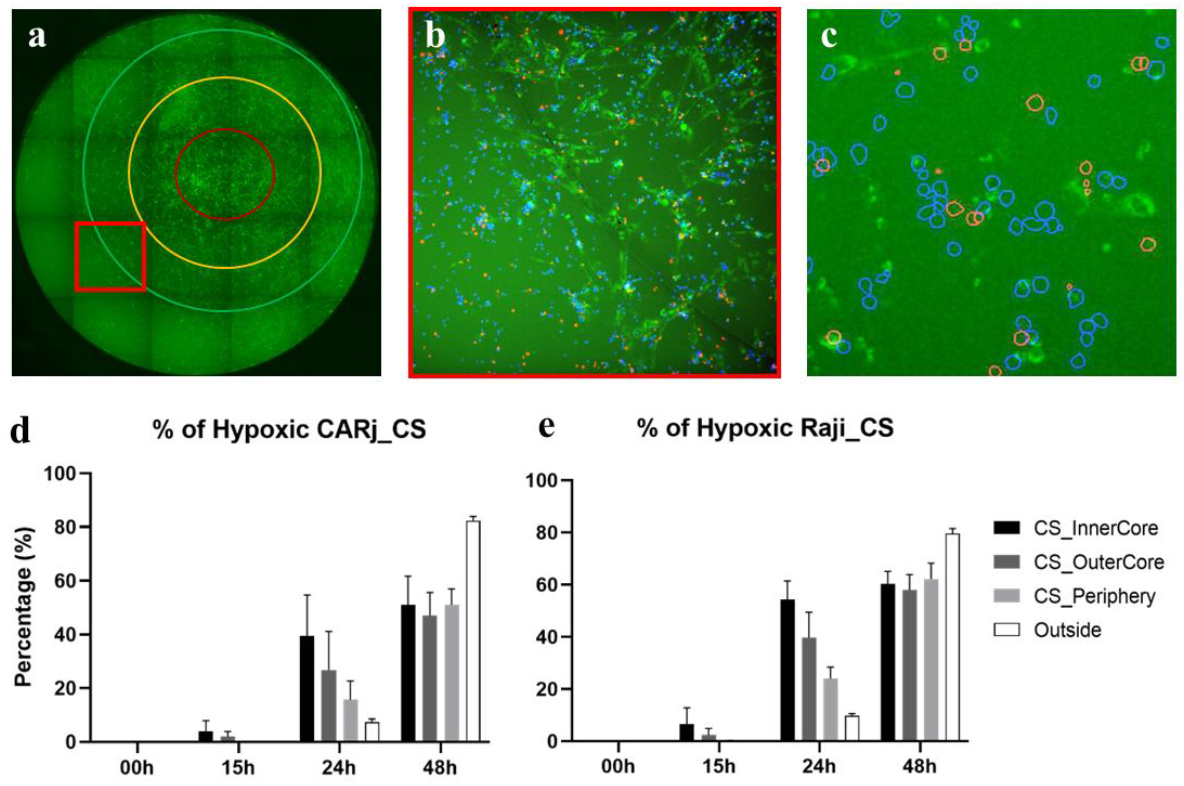
Spatial quantification of hypoxic cells in the CSH model. (a) Composite well image acquired in the FITC channel with overlaid spatial zones (InnerCore, OuterCore, Periphery, and Outside). (b) The red inset highlights one acquisition field, showing Raji cells (red), CAR-Jurkat cells (blue), and hypoxia labeling (green). (c) High-magnification micrograph displaying segmented Raji (red) and CAR-Jurkat (blue) cells overlaid onto the FITC hypoxia channel, enabling direct association of hypoxic signal with individual cells. (d– e) Quantitative analysis of the percentage of hypoxic CAR-Jurkat (d) and Raji (e) cells over time (0–48 h) across spatial zones. The data reveal the progressive development of hypoxia beneath the coverslip, first emerging within the InnerCore at ∼15 h and displaying a decreasing spatial gradient from InnerCore to Outside by 24 h, thereby validating the computational segmentation and oxygen-gradient mapping strategy used for high-content screening. Bars represent mean ± SD from three independent experiments (p < 0.05; two-way ANOVA with Šídák correction).

**Figure 4.**
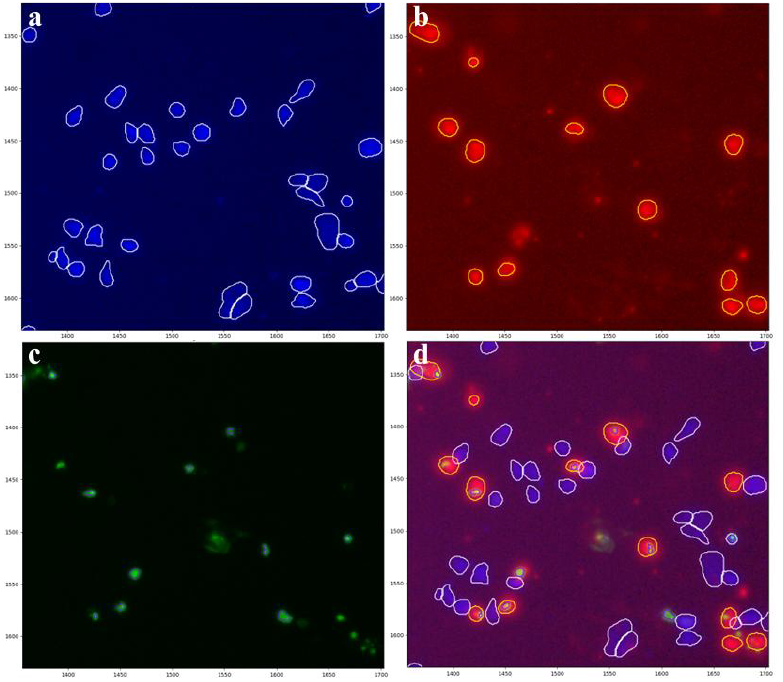
Automated segmentation using CellProfiler pipelines. Representative images illustrating the segmentation workflow applied to the Coverslip Hypoxia High-Content Screening (CH-HCS) assay. (a) Raw fluorescence image acquired with the DAPI filter cube, showing anti-CD19 CAR-T cells stained with CellTrace Violet (CTV). Automated segmentation (outlined in white) generated using RunCellpose, an AI-based GPU-powered CellProfiler plugin module. (b) Fluorescence image acquired with the Cy5 filter cube, corresponding to CD19^+^ Raji tumor cells pre-labeled with Cell Proliferation eFluor 670. Segmentation of Raji cells is outlined in yellow. (c) Fluorescence image acquired with the FITC filter cube, indicating Sytox Green - positive cells (death marker). Automated segmentation of dead cells is outlined in purple. (d) Final overlay of segmented objects combining the DAPI, Cy5 and FITC channels. This panel shows Raji cells (yellow outlines), CAR-T cells (white outlines) and dead cells (purple outlines) highlighting the attribution of dead cells (green) to the specific population. These images collectively demonstrate the capacity of the CH-HCS analysis pipeline to distinguish multiple cell populations and viability states within a single field, enabling region-resolved quantification of CAR-T cytotoxicity and hypoxia-dependent effects across the tumor–stroma interface.

### 2.9 Data handling, statistics, and replication

Per-ROI single-cell and per-well summaries were exported from CellProfiler, processed in KNIME Analytics Platform, and analyzed in GraphPad Prism (v9; v8 where specified in figure legends). Unless otherwise indicated, data are presented as mean ± SEM from three independent experiments with three technical replicates (wells) per condition. For statistical analyses, per-well aggregated values were used as the unit of analysis. Group comparisons were performed using ordinary two-way ANOVA with Šídák’s multiple-comparisons test; significance thresholds and exact *p* values are reported in the figure legends.

## 3 Results

Quantitative analysis of cell death in the 2D Coverslip Hypoxia (CSH) model revealed clear spatial and functional differences between CAR-T–mediated cytotoxicity and control conditions. In co-cultures of CD19^+^ Raji cells with anti-CD19 CAR-T lymphocytes (Figure 5a,e), tumor cell death increased significantly over 24 hours, reaching up to ∼60 % in normoxic regions (Outside) and ∼40–50 % in peripheral/transitional zones, while remaining below 10 % within hypoxic InnerCore and OuterCore regions (two-way ANOVA, *p* values ranging from *< 0.01 to < 0.0001*). In contrast, co-cultures with non-transduced T cells (Figure 5b,f) produced minimal Raji death (< 15 %) across all oxygen-defined regions, confirming antigen specificity. For CD19^−^ K562 cells (Figure 5c,d,g,h), negligible cytotoxicity was observed in both CAR-T and control T cell conditions (< 10–15 %), indicating the absence of off-target effects. Interestingly, effector cell viability exhibited the inverse trend: both CAR-T and non-transduced T cells displayed reduced cell death under hypoxic conditions (Inner/Outer Core) compared with normoxic regions (*p < 0.05–0.001*), suggesting that low-oxygen niches confer a relative survival advantage. Collectively, these findings demonstrate that the CH-HCS assay captures spatially resolved immune-effector dynamics, revealing that hypoxia selectively impairs CAR-T cytotoxicity while concurrently protecting both tumor and effector cells—recapitulating critical features of the tumor microenvironment.

**Figure 5.**
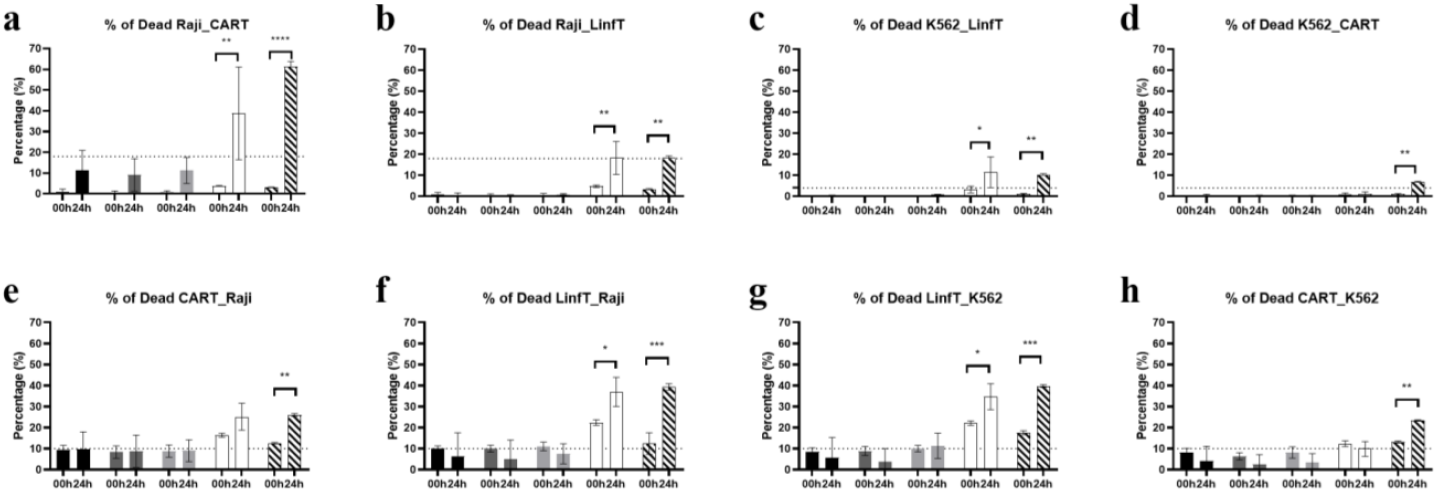
Percentage of tumor and effector cell death in 2D Coverslip Hypoxia co-cultures. Bar graphs show the percentage of dead cells (Y axis) in distinct oxygen-defined regions of the CSH assay—InnerCore (severe hypoxia), OuterCore (moderate hypoxia), Periphery (transition), and Outside (normoxia)—at 0 h and 24 h. (a,e) Raji + CAR-T; (b,f) Raji + T cells; (c,g) K562 + CAR-T; (d,h) K562 + T cells. CAR-T–mediated cytotoxicity toward CD19^+^ Raji cells was significantly higher in normoxic and peripheral regions, while hypoxic cores exhibited minimal killing. Non-transduced T cells and all K562 co-cultures showed negligible cytotoxicity, confirming antigen specificity. Effector (T cell and CAR-T) death was greater in normoxic regions, decreasing toward hypoxia, consistent with oxygen-dependent metabolic stress. Bars represent mean ± SD from three independent experiments; significance assessed by two-way ANOVA with Šídák’s post-test (*p < 0.05, **p < 0.01, ***p < 0.001, \*\*\**p < 0.0001*).

**Figure 6.**
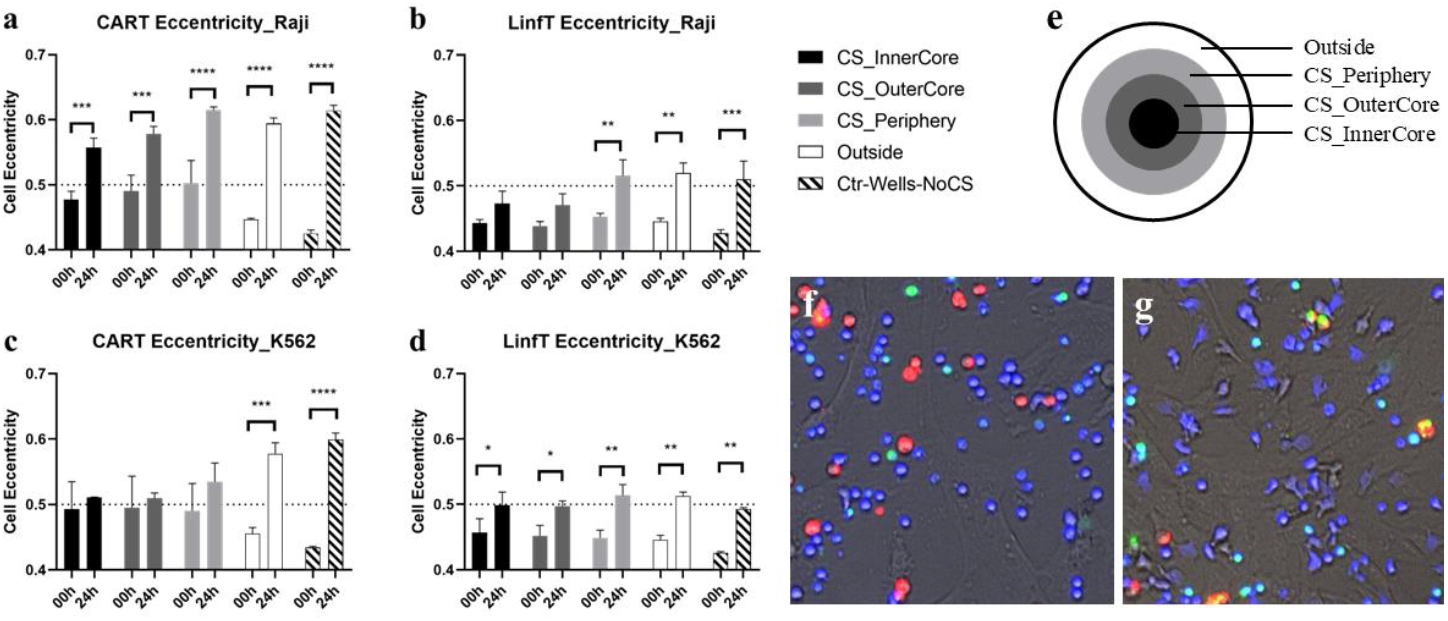
Morphometric analysis of T lymphocytes and anti-CD19 CAR-T cells in the 2D Coverslip Hypoxia co-culture model. Bar graphs represent mean ± SD cell eccentricity (Y axis; 0 = circular, 1 = elongated) measured at 0 h and 24 h across oxygen-defined regions of the CSH model: **CS_InnerCore, CS_OuterCore, CS_Periphery, Outside**, and control wells without coverslip (**Ctr-Wells-NoCS**). (a) CAR-T + Raji; (b) T cells + Raji; (c) CAR-T + K562; (d) T cells + K562. (e) Schematic representation of the spatial composition of the regions of interest (f) Representative 0 h acquisition showing rounded resting effector cells. (g) Representative 24 h acquisition showing elongated, activated CAR-T cells interacting with tumor targets. Statistical analysis: two-way ANOVA with Šídák’s multiple-comparison test (*p < 0.05, **p < 0.01, ***p < 0.001, \*\*\**p < 0.0001*). Data are presented as mean ± SEM from three independent biological experiments.

Morphometric analysis of T lymphocytes and anti-CD19 CAR-T cells co-cultured with Raji (CD19^+^) or K562 (CD19^−^) tumor cells revealed oxygen- and antigen-dependent changes in cell shape dynamics. Eccentricity, a quantitative measure of cell elongation (0 = circular; 1 = elongated), was used as a proxy for migratory or activation status. In CAR-T + Raji co-cultures (Figure 6a), mean eccentricity increased significantly after 24 h across all oxygen-defined regions, reaching values close to 0.6 in normoxic and peripheral zones, consistent with enhanced motility and cytotoxic engagement. Unmodified T cells co-cultured with Raji (Figure 6b) displayed only modest eccentricity increases (0.45–0.5), suggesting limited activation. In K562 co-cultures (Figures 6c,d), both CAR-T and control T cells maintained low eccentricity under hypoxic conditions, with small but significant increases in normoxic areas, reflecting baseline migratory behavior without antigen-specific activation. Representative fluorescence micrographs (Figures 6e,f) illustrate the morphological transition of CAR-T cells from rounded, resting profiles at 0 h to elongated and polarized shapes at 24 h, consistent with activation and target engagement. Together, these data demonstrate that CAR-T motility and activation are strongly influenced by both antigen recognition and local oxygen availability, confirming that hypoxia constrains effector morphology and function in the CH-HCS assay.

To investigate whether inhibition of the adenosinergic signaling axis could enhance CAR-T cytotoxicity under hypoxic conditions, we treated 2D CAR-T/Raji co-cultures with the selective A_2_A receptor antagonist Ciforadenant and compared them with DMSO controls. Quantitative analysis revealed that Ciforadenant consistently increased the percentage of dead tumor cells across all spatial regions of the Coverslip Hypoxia (CSH) model throughout the 40 h time course. The effect was most pronounced in the hypoxic InnerCore and OuterCore regions, where CAR-T cytotoxicity is typically attenuated. In these zones, cell death reached approximately 60–70 % by 40 h in the Ciforadenant-treated group versus 35–45 % in DMSO controls. Two-way ANOVA indicated a significant interaction between time and oxygen level (p < 0.01), reflecting distinct temporal patterns of cytotoxicity across oxygen gradients, with both time (p < 0.0001) and oxygen level (p < 0.1) contributing as main effects (Figure 7). These findings support the hypothesis that blocking A_2_A receptor signaling partially restores CAR-T effector function within oxygen-restricted microenvironments, alleviating the immunosuppressive influence of extracellular adenosine accumulation in hypoxia.

**Figure 7.**
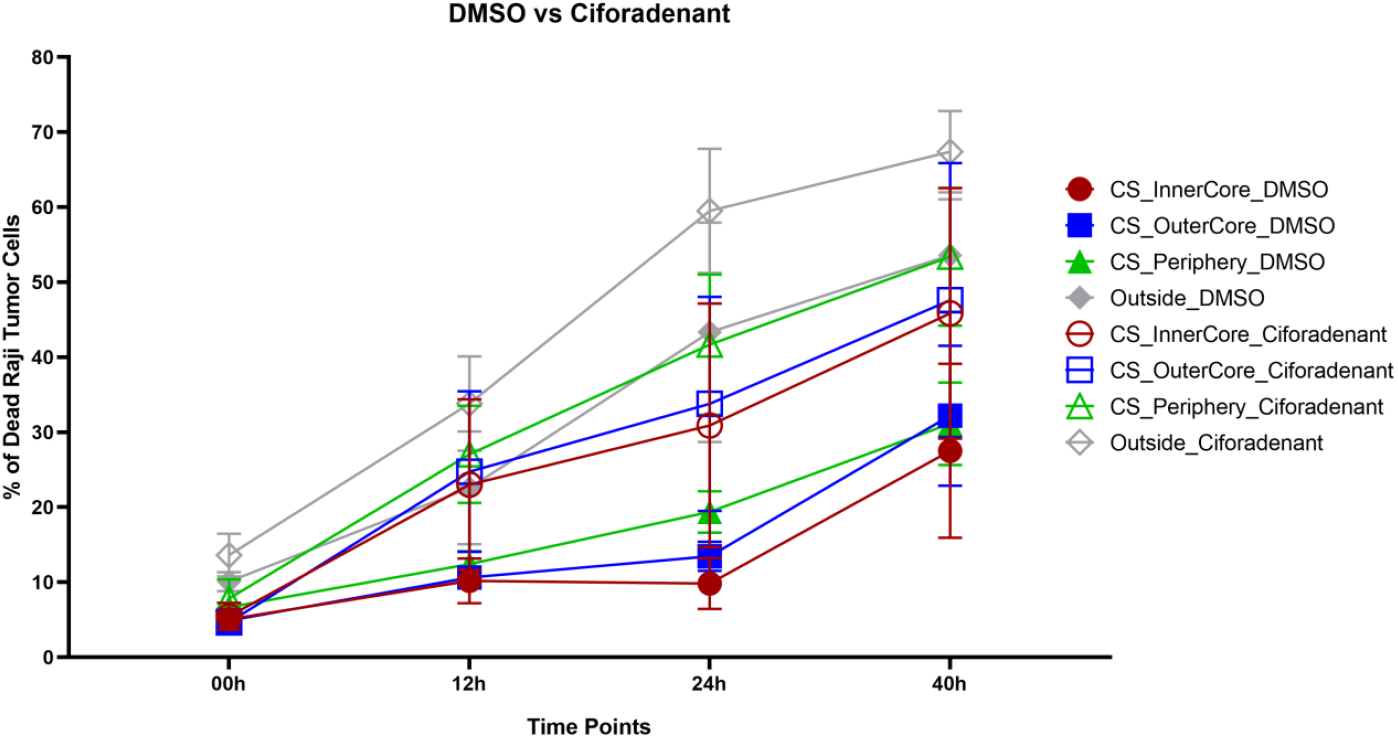
Effect of A_2_A receptor blockade on CAR-T cytotoxicity in a 2D co-culture model. Percentage of dead CD19^+^ Raji tumor cells co-cultured with anti-CD19 CAR-T cells under different oxygen-defined regions of the Coverslip Hypoxia system (InnerCore, OuterCore, Periphery) over time (0, 12, 24, 40 h). The curves compare treatment with the A_2_A receptor antagonist Ciforadenant (open symbols) versus vehicle control (0.1 % DMSO, solid symbols). A progressive increase in tumor-cell death was observed in all regions, with greater cytotoxicity in Ciforadenant-treated wells, particularly within hypoxic cores. Statistical analysis by two-way ANOVA showed significant interaction of time × oxygen level (p < 0.01), time effect (p < 0.0001), and oxygen-level effect (p < 0.1). Data represent mean ± SEM from three independent experiments.

## 4 Discussion

The Coverslip Hypoxia High-Content Screening (CH-HCS) platform presented here provides a quantitative, spatially resolved approach to assess how oxygen gradients modulate CAR-T activity, viability, and morphodynamics within a simplified two-dimensional tumor–stroma interface. By integrating controlled hypoxia with high-content fluorescence imaging and automated segmentation, this method bridges a critical gap between conventional 2D co-cultures and complex 3D models, offering the throughput necessary for systematic pharmacological or genetic screening.

Our results demonstrate that local oxygen availability exerts profound effects on CAR-T cytotoxicity and behavior. In the CH-HCS assay, anti-CD19 CAR-T cells displayed robust and antigen-specific cytotoxicity against CD19^+^ Raji cells in normoxic and peripheral regions, yet their killing capacity was markedly reduced within hypoxic InnerCore and OuterCore regions. This spatial heterogeneity mirrors clinical and in-vivo findings showing that T cells infiltrating hypoxic tumor niches exhibit impaired effector function due to metabolic restriction and transcriptional reprogramming. Reduced cytotoxicity in low-oxygen zones is likely associated with diminished glycolytic flux and ATP generation, reduced granzyme B and IFN-γ secretion, and the accumulation of extracellular adenosine and lactate—factors known to suppress T-cell activation through A2A/A2B receptor signaling.

Conversely, both CAR-T and unmodified T cells exhibited increased survival under hypoxic conditions, consistent with reports that oxygen deprivation reduces activation-induced cell death and promotes quiescence. This survival advantage may reflect a metabolic shift toward oxidative quiescence or enhanced resistance to reactive oxygen species. Together, these results highlight a dual influence of hypoxia—protecting effector cells from apoptosis while simultaneously dampening their cytotoxic output. Such spatially opposed outcomes underscore the importance of considering microenvironmental context when evaluating therapeutic potency.

Morphometric analysis further revealed that CAR-T activation and motility, inferred from cell eccentricity, were strongly dependent on both antigen presence and oxygen tension. CAR-T cells co-cultured with CD19^+^ targets exhibited pronounced elongation and polarization in normoxic regions, characteristic of active immunological synapse formation and directional migration. Under hypoxia, CAR-T cells remained round and static, indicating suppressed cytoskeletal remodeling and reduced migratory and surveillance activity. These quantitative imaging metrics complement traditional cytotoxicity assays by capturing dynamic, non-lethal functional states that precede target killing.

The simplicity and adaptability of the CH-HCS platform confer several advantages over existing hypoxia models. The radial oxygen gradient created by the coverslip provides co-localized hypoxic and normoxic microdomains within a single well, enabling internal normalization and direct comparative analysis. Combined with automated image segmentation and open-source analytics (CellProfiler and KNIME), this configuration supports unbiased, high-throughput assessment of spatially heterogeneous responses. Importantly, the model is compatible with primary tumor cells, engineered T cells, and stromal components, facilitating preclinical exploration of combination therapies—such as A2A/A2B receptor antagonists, HIF pathway inhibitors, or metabolic reprogramming strategies—to restore CAR-T efficacy in hypoxic tumors.

The addition of Ciforadenant to the 2D Coverslip Hypoxia model demonstrates that pharmacologic inhibition of A_2_A receptor signaling can partially reverse hypoxia-induced suppression of CAR-T function. This finding reinforces the mechanistic role of adenosine accumulation and A_2_A receptor activation as key mediators of effector-cell dysfunction in oxygen-deprived tumor niches. By preventing A_2_A-driven cAMP elevation and downstream PKA signaling, Ciforadenant likely preserved T-cell receptor signaling and cytolytic capacity, particularly within the InnerCore and OuterCore regions where hypoxia was most severe. The observed restoration of tumor-cell killing parallels in vivo reports. In a phase I clinical trial of 68 patients with renal cell carcinoma—including individuals refractory to PD-1/PD-L1 checkpoint blockade—Ciforadenant, alone or combined with anti–PD-L1 therapy, demonstrated durable clinical benefits associated with increased intratumoral CD8^+^ T-cell infiltration and a broadened circulating T-cell repertoire^(*118*)^. Together, these results highlight the CH-HCS assay’s capacity to quantify drug efficacy across physiological oxygen gradients and support the therapeutic potential of adenosinergic pathway inhibitors to improve CAR-T performance in the metabolically hostile microenvironments typical of solid tumors.

Classical methods to evaluate the functional activity of cytotoxic lymphocytes, including CAR-T cells, were historically designed to measure either activation of the effector cell or death of the target cell population, but each approach presents intrinsic limitations in discriminating specific from nonspecific effects and in capturing spatiotemporal dynamics. Functional assays sought to quantify cytotoxicity toward target cells, generally use population-based measurements of membrane integrity. For instance, the traditional ^51^Cr-release assay ^(*119*)^ remains a benchmark but is limited by radioactive handling, endpoint readout, and specifically suited only to quantify chromium release from the dead cell population that was subjected to the incorporation of ^51^Cr. The LDH-release assay, which quantifies (via colorimetric or fluorometric readouts) the activity of the cytosolic enzyme lactate dehydrogenase released from damaged cells eliminates the use of radioactivity, but depends on the assumption that effector cells remain >95% viable, as their own death also releases LDH, thus contributing to the signal ^(*120*)^. Other target-specific assays, similar to the ^51^Cr release assay, such as the fluorometric Calcein-AM release assay, which uses target cells labeled intracellularly with a fluorescent dye ^(*121*)^, or luminescent systems, based on target cell lines genetically modified to express luciferase in the cytosol ^(*122-124*)^, improved sensitivity and reproducibility, but remain bulk measurements—providing a single average value that cannot distinguish selective killing of specific subpopulations within mixed cultures. Moreover, these assays quantify loss of target-cell integrity but not effector persistence, proliferation, or death, all of which are critical determinants of CAR-T efficacy.

More refined “label-free” or impedance-based assays ^(*125, 126*)^ enable real-time monitoring of adherent target-cell detachment, yet this metric is only an indirect proxy for cytotoxicity and poorly suited for non-adherent hematologic tumor models. To overcome such limitations, flow-cytometry-based cytotoxicity assays label target cells with CFSE or equivalent dyes and quantify their disappearance in mixed cultures^(*127*)^. Variations such as the VITAL assay ^(*128*)^ allow multiplexed assessment of multiple target populations stained at distinct fluorescence intensities, thereby controlling for off-target or antigen-independent effects. However, flow cytometry disrupts the physical organization of co-cultures and precludes kinetic observation, offering only static snapshots of cell death. Imaging-based cytometry further advanced quantification by directly counting live (GFP^High^) and dead (GFP^Low^) target cells ^(*129*)^, but, relying on uniform 2D cultures lacking the stromal component and the gradients of oxygen and metabolites characteristic of the tumor microenvironment (TME).

To recreate such gradients, 3D multicellular tumor spheroid models have been adopted ^(*130-133*)^. Tumor-stroma spheroids are valuable for studying immune-cell infiltration and tumor killing within tissue-like architecture ^(*106-110*)^. Yet, because of their large size (>500 µm), spheroids present optical and analytical challenges: imaging penetration is limited, single-cell segmentation is difficult, and kinetic measurements are impractical or limited. Confocal microscopy of immune-cell infiltration, combined with hypoxia markers such as pimonidazole ^(*105*)^ provides static cross-sections but not live, high-throughput quantification of T-cell function.

Our Coverslip Hypoxia High-Content Screening (CH-HCS) assay was designed to bridge the gap between the accessibility of 2D co-cultures and the physiological relevance of 3D models. By placing a 5 mm glass coverslip over tumor–stroma co-cultures, the system generates reproducible radial oxygen and metabolic gradients within a single well, creating regions analogous to the normoxic periphery and hypoxic core of solid tumors. Using automated CellProfiler pipelines, the assay defines spatial Regions of Interest (ROIs)—InnerCore, OuterCore, Periphery, and Outside— allowing independent quantification of each compartment. Unlike traditional bulk assays, CH-HCS provides single-cell, spatially resolved measurements of both tumor and effector populations across time, under identical conditions. Because tumor (e.g., Raji) and effector (CAR-T) cells are pre-labeled with distinct fluorophores, their proliferation, death, and relative distribution can be followed in real time through live imaging and analyzed in high throughput.

Functionally, our approach enables parallel quantification of CAR-T cytotoxicity, effector survival, and proliferation dynamics within microenvironments of different oxygen availability— essentially yielding a “cross-sectional” view of a tumor spheroid while retaining the throughput and optical accessibility of 2D cultures. The method overcomes major sources of confusion in classical cytotoxicity assays, as target-cell and effector-cell fates are measured simultaneously rather than inferred indirectly from population averages. Furthermore, by maintaining stable hypoxia gradients and permitting time-lapse analysis, CH-HCS reveals the evolving balance between CAR-T activity and tumor resistance mechanisms under conditions that more closely mimic the tumor–stroma interface in vivo. This spatially and temporally resolved design thus represents a powerful evolution of functional CAR-T assays, integrating activation, cytotoxicity, and metabolic context within a unified, quantitative framework.

Of note, combining the CH-HCS approach with reporter cell lines carrying fluorescent reporters under NF-κB, NFAT, or AP-1–responsive promoters ^(*134-143*)^, would enable automated readouts of T-cell activation, co-stimulation, or inhibition, in addition to target-cell fate, spatial dynamics, and effector viability.

## 5 Conclusion

This work establishes the Coverslip Hypoxia High-Content Screening (CH-HCS) platform as a robust and quantitative approach to model tumor–immune interactions across physiologically relevant oxygen gradients. By integrating high-resolution imaging, automated spatial segmentation, and region-specific analysis, the system enables simultaneous evaluation of CAR-T cytotoxicity, survival, and morphodynamics under normoxic and hypoxic conditions within a single well. The addition of the A_2_A receptor antagonist Ciforadenant validated the model’s sensitivity to pharmacologic modulation, demonstrating that adenosinergic blockade can mitigate hypoxia-driven immune suppression. Collectively, the CH-HCS platform provides a scalable and reproducible tool for dissecting microenvironmental constraints on CAR-T therapy and for preclinical screening of metabolic or immunomodulatory strategies aimed at restoring effector function in solid tumors.

## 6 Acknowledgments

We gratefully acknowledge Dr. Amanda Mizukami Martins (Production Manager at NUTERA – Advanced Cell Therapy Center, Hemocentro Ribeirão Preto) for kindly providing the anti-CD19 CAR-T cell line and T lymphocytes used in this study. As well, we thank the Cancer Proteomics Research Laboratory (Prof. Dr. Vitor Marcel Faça) for generously donating the anti-CD19 CAR-Jurkat cell line used in this work.

This study was funded by the São Paulo Research Foundation (FAPESP), Brasil, process #2022/12856-6; the Brazilian Federal Agency for Support and Evaluation of Graduate Education (CAPES); and the National Council for Scientific and Technological Development (CNPQ).

---

**Supplementary Figure S1.**
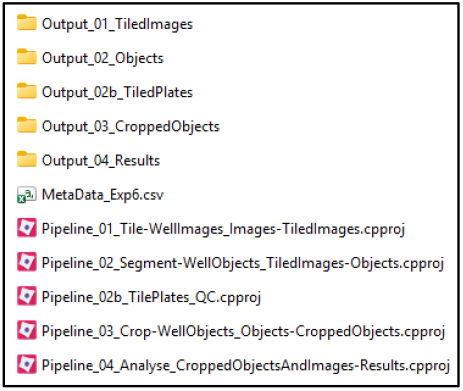
Directory structure and workflow organization of the *CellProfiler* project for Coverslip Hypoxia High-Content Screening (CH-HCS) analysis. The folder hierarchy contains sequentially executed pipelines and their corresponding output directories: (i) Pipeline_01 – image tiling and metadata extraction (*Output_01_TiledImages*); (ii) Pipeline_02 – object segmentation (*Output_02_Objects*); (iii) Pipeline_02b – plate-level quality control (*Output_02b_TiledPlates*); (iv) Pipeline_03 – cropping and region-of-interest isolation (*Output_03_CroppedObjects*); and (v) Pipeline_04 – final quantification and data export (*Output_04_Results*). The *MetaData_Exp6.csv* file contains experiment-wide identifiers used for automated grouping and tracking across the analysis stages.

**Supplementary Figure S2.**
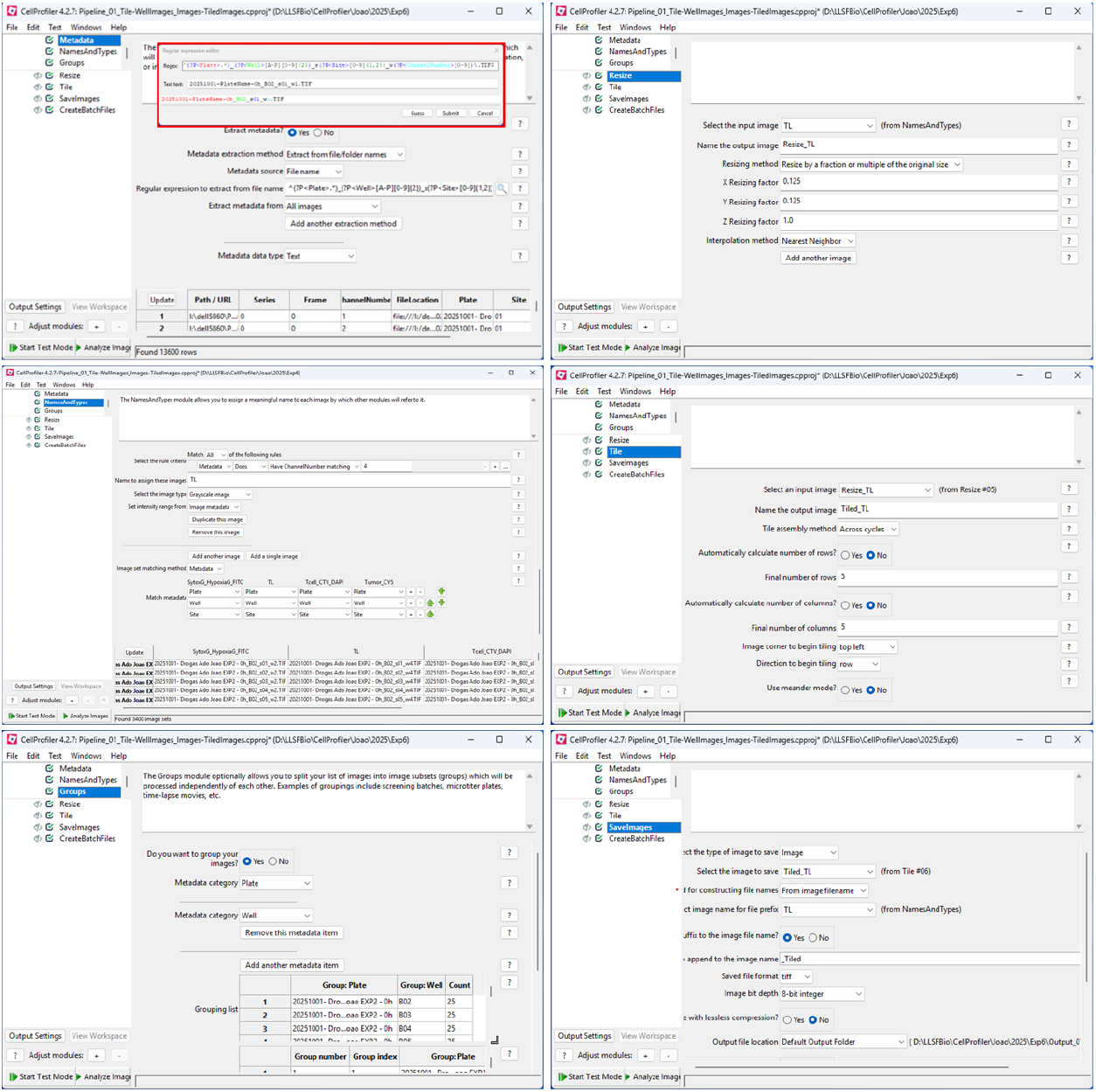
Overview of the *CellProfiler* pipeline used for image preprocessing and well-stitching for CH-HCS. Representative screenshots of the *CellProfiler 4.2.7* interface show the sequential configuration of modules used for image processing in the pipeline “Pipeline_01_Tile-WellImages_Images-TiledImages.cpproj”. The pipeline included the following steps: (i) **Metadata** module, defining regular expressions and metadata features for plate and well identifiers; (ii) **Resize** module, reducing image resolution to 12.5% of the original for computational efficiency; (iii) **Tile** module, assembling 6×5 image grids corresponding to each well into full-well composite images; (iv) **NamesAndTypes** module, defining input file selection and image type; (v) **Groups** module, defining metadata-based image grouping by plate and well identifiers; and (vi) **SaveImages** module, exporting tiled TIFF files for downstream segmentation and analysis.

**Supplementary Figure S3.**
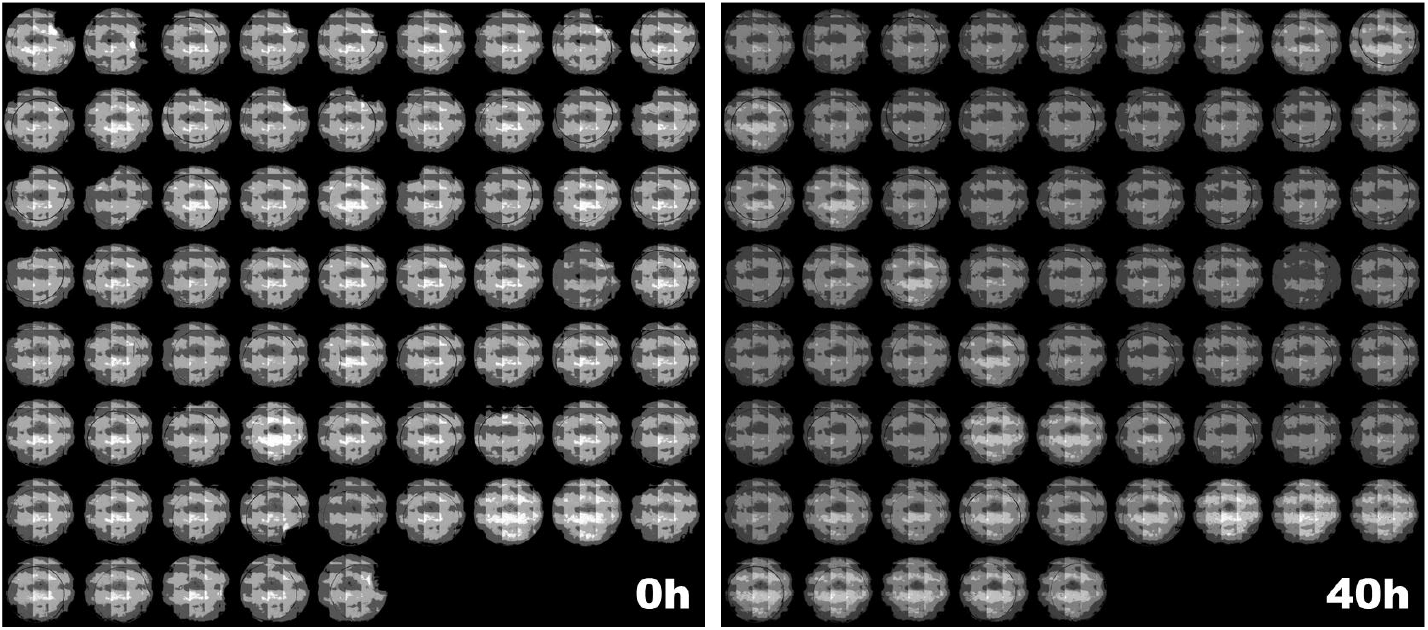
Quality control of coverslip positioning across time points using the “Pipeline_02b_TilePlates_QC.cpproj”. Representative stitched transmitted-light images of 96-well plates at the beginning (0 h) and end (40 h) of the experiment. The *CellProfiler* pipeline “Pipeline_02b_TilePlates_QC.cpproj” was used to evaluate the stability of coverslip positioning during long-term imaging in the Coverslip Hypoxia (CSH) assay. Each well image was tiled and aligned to compare coverslip placement between time points, ensuring consistent spatial reference for oxygen-gradient quantification. Coverslips must remain fixed throughout the experiment to maintain stable hypoxia-defined Regions of Interest (ROIs). Once the plate passes this quality control step—showing no significant coverslip displacement—only the images from a single time point (typically 0 h) are processed in the segmentation pipeline “Pipeline_02_Segment-WellObjects_TiledImages-Objects.cpproj” to define the spatial ROIs used in subsequent high-content analyses.

**Supplementary Figure S4.**
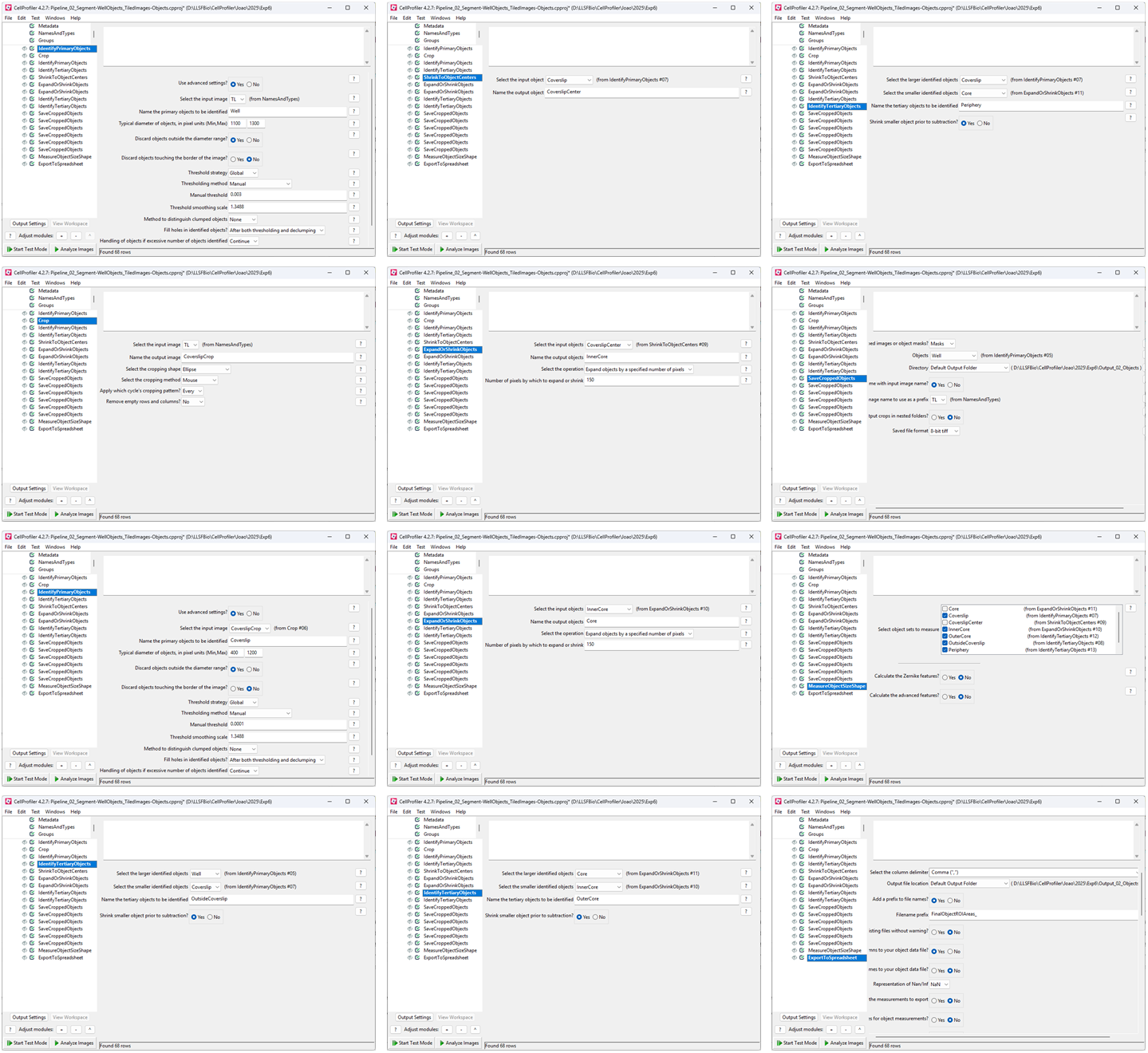
Overview of the *CellProfiler* pipeline used for defining spatial Regions of Interest (ROIs) in each well of the Coverslip Hypoxia CH-HCS model. The *CellProfiler* pipeline (*Pipeline_02_Segment-WellObjects_TiledImages-Objects.cpproj*) was designed to automatically delineate spatial compartments corresponding to different oxygenation zones within each well. The well ROI is segmented automatically from transmitted light images, as the use of black-walled plates provides high contrast between the illuminated well area and the background. The coverslip ROI is manually defined for each well, serving as the central hypoxic region; however, this step can be automated in future applications using coverslips that block transmitted light. All remaining ROIs—InnerCore, OuterCore, Periphery, and Outside—are generated computationally and hierarchically related to each other by Boolean operations (object subtraction and expansion) within the same pipeline. This automated spatial segmentation framework enables consistent definition of oxygen-dependent regions across multiple wells, providing the foundation for quantitative single-cell mapping of hypoxia gradients in high-content screening assays.

**Supplementary Figure S5.**
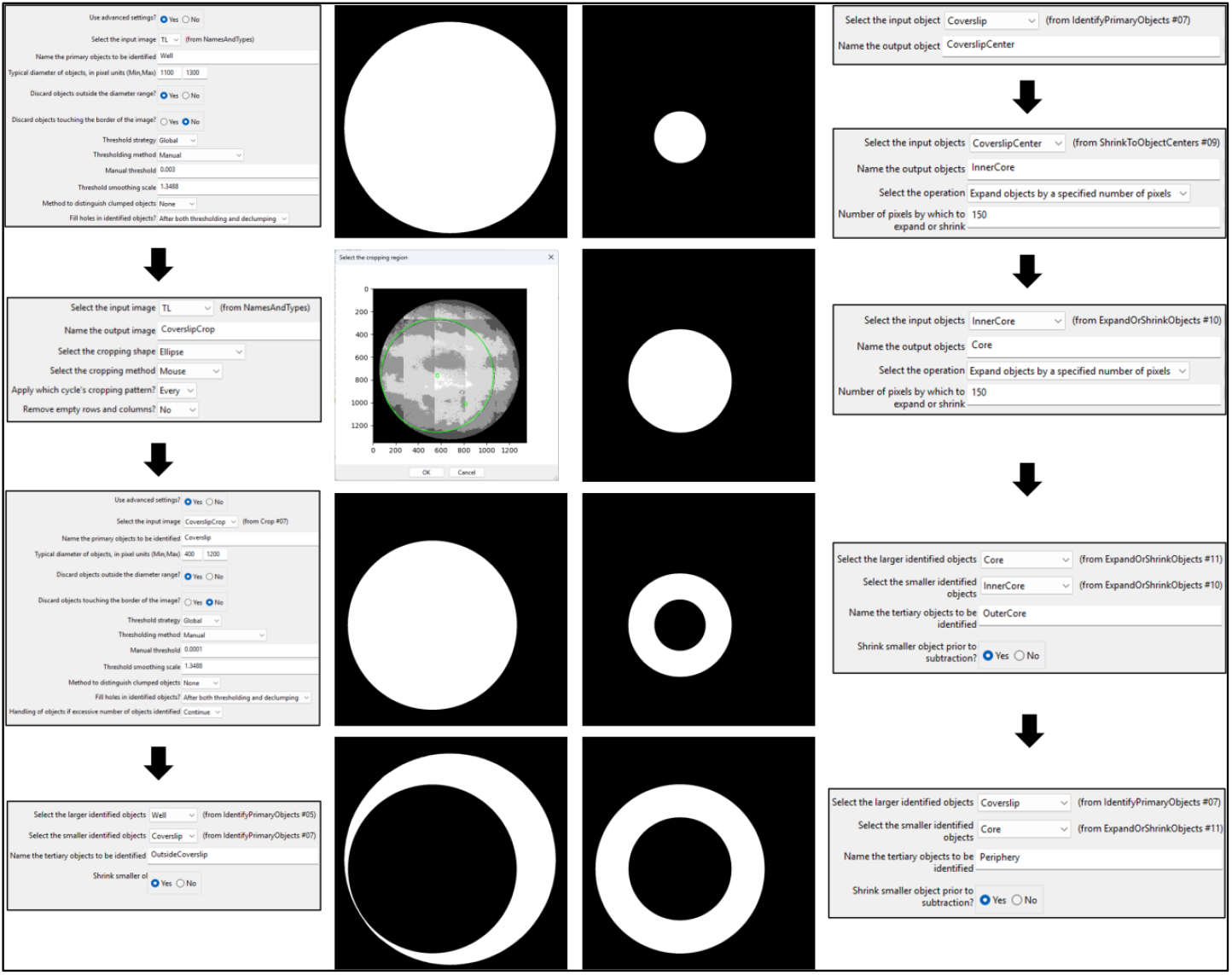
Representative Regions of Interest (ROIs) in the CSH model obtained using CellProfiler. The pipeline illustrates the sequential steps for defining spatial compartments within each well based on transmitted light images. The well ROI is automatically segmented due to the contrast provided by black-walled plates. The coverslip region is manually cropped using an elliptical selection tool, defining the hypoxia-inducing zone. From this region, the InnerCore (severe hypoxia) is generated by shrinking the coverslip center, while the Core and OuterCore regions are produced by expanding the InnerCore by fixed pixel distances (150 px per step). The Periphery region, representing the hypoxia-to-normoxia transition zone, is automatically defined by subtracting the Core from the Coverslip, and the OutsideCoverslip (normoxic region) is created by subtracting the Coverslip from the total well area. Together, these hierarchical operations generate spatially related ROIs that accurately represent the radial oxygen gradient for quantitative mapping of cellular responses under the coverslip.

**Supplementary Figure S6.**
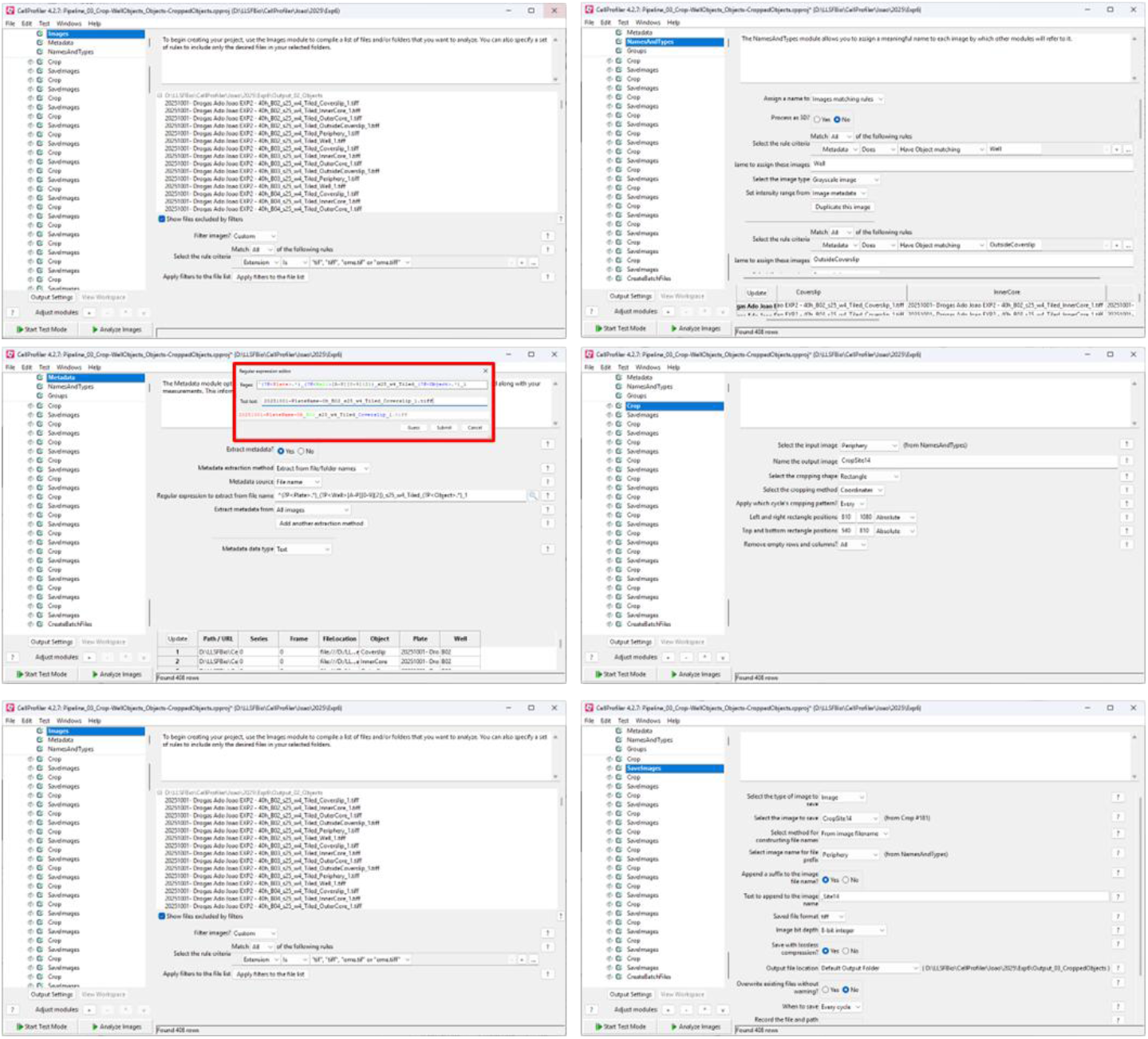
Automated cropping workflow pipeline. The *CellProfiler* pipeline “Pipeline_03_Crop-WellObjects_Objects-CroppedObjects.cpproj” performs automated cropping of the whole objects obtained by segmentation of ROIs from the full-well composite images (a 5×5 tiled mosaic) into 25 smaller rectangular sub-images, each corresponding to one acquisition site from the original tiled image. The workflow begins with the Metadata and NamesAndTypes modules to import image metadata and assign channel identities, followed by the Groups module to organize wells for batch processing. The Crop module then applies spatial coordinates to divide the full-well object image into 25 equal fields, generating one cropped object per site while preserving spatial correspondence to the original well layout. Finally, cropped ROIs are exported as individual TIFF files via the SaveImages module. These cropped objects are subsequently used in the next pipeline (“Pipeline_04_Analyse_CroppedObjectsAndImages-Results.cpproj”), where they are resized back to their original dimensions and overlaid onto fluorescence-channel images, allowing segmented CTV-stained CAR-T cells (from the DAPI image) and CP-670–labeled Raji cells (from the Cy5 channel) to be mapped to their precise spatial ROI within each well. Note the inset in red with the regular expression that captures the object name added as a suffix in the previous pipeline.

**Supplementary Figure S7a.**
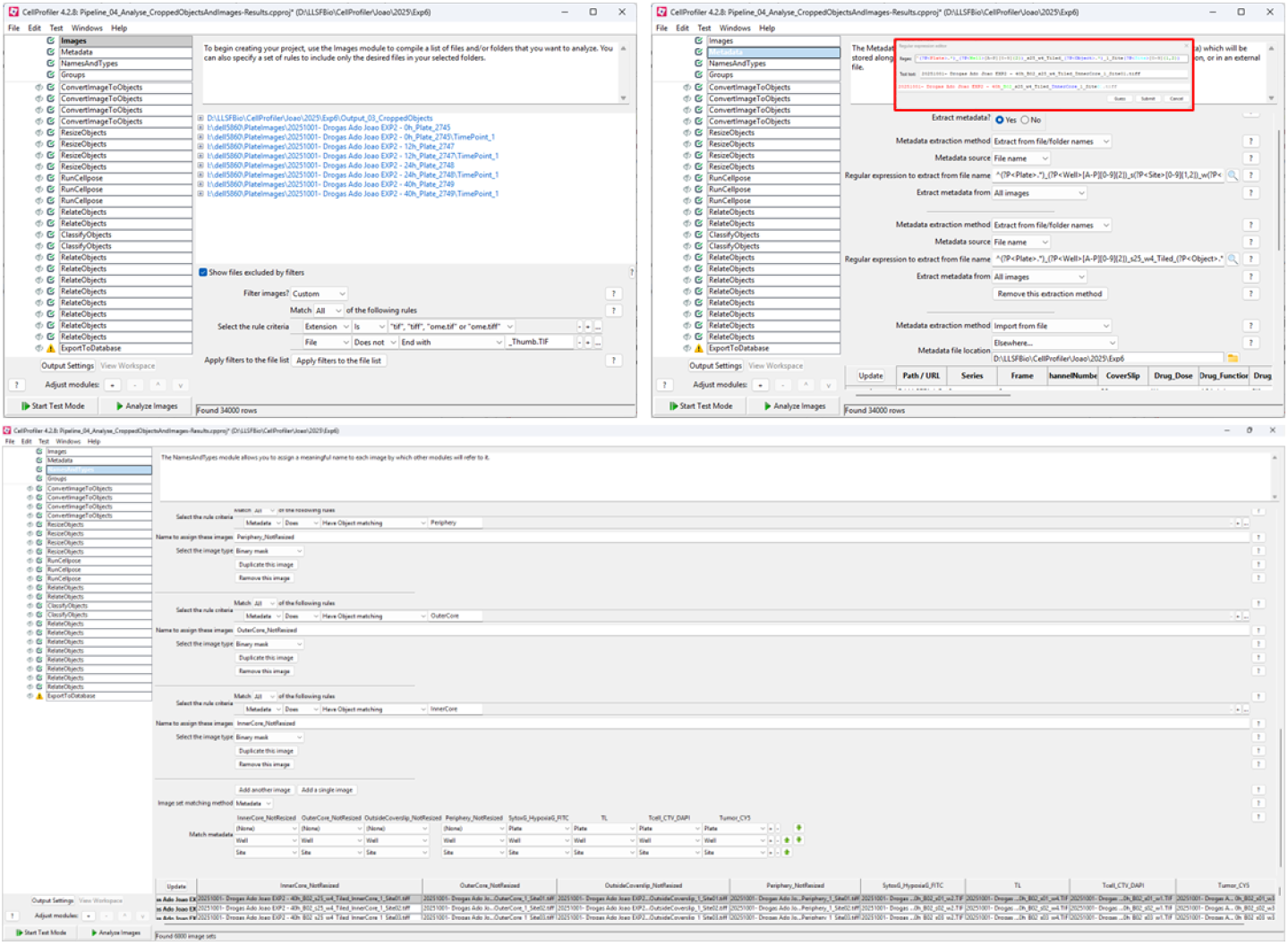
a. Structure and workflow of the pipeline “Pipeline_04_Analyse_CroppedObjectsAndImages-Results.cpproj”. Composite image showing the main modules of the *CellProfiler 4.2.8* pipeline used for single-cell quantification and spatial mapping of CAR-T and tumor cell populations across oxygen-defined regions of the Coverslip Hypoxia (CSH) model. The pipeline begins by loading all fluorescence channels (DAPI, FITC, Cy5, transmitted light) at full resolution and the cropped ROI objects corresponding to each of the 25 tiled acquisition sites per well. In the **Metadata** module, regular expressions extract *Plate, Well*, and *Site* information from image filenames, while cropped object filenames provide *Object* identifiers, as well as *Well*, and *Site* information. These metadata fields are linked with the experimental CSV file, ensuring that each image is associated with its biological and experimental condition (drug, dose, replicate, etc). The **NamesAndTypes** module assigns names to each fluorescence channel and cropped object and aligns them by plate, well, and site, while objects are aligned only by well and site, allowing consistent reuse of segmented ROI masks across different timepoints.

**Supplementary Figure S7b.**
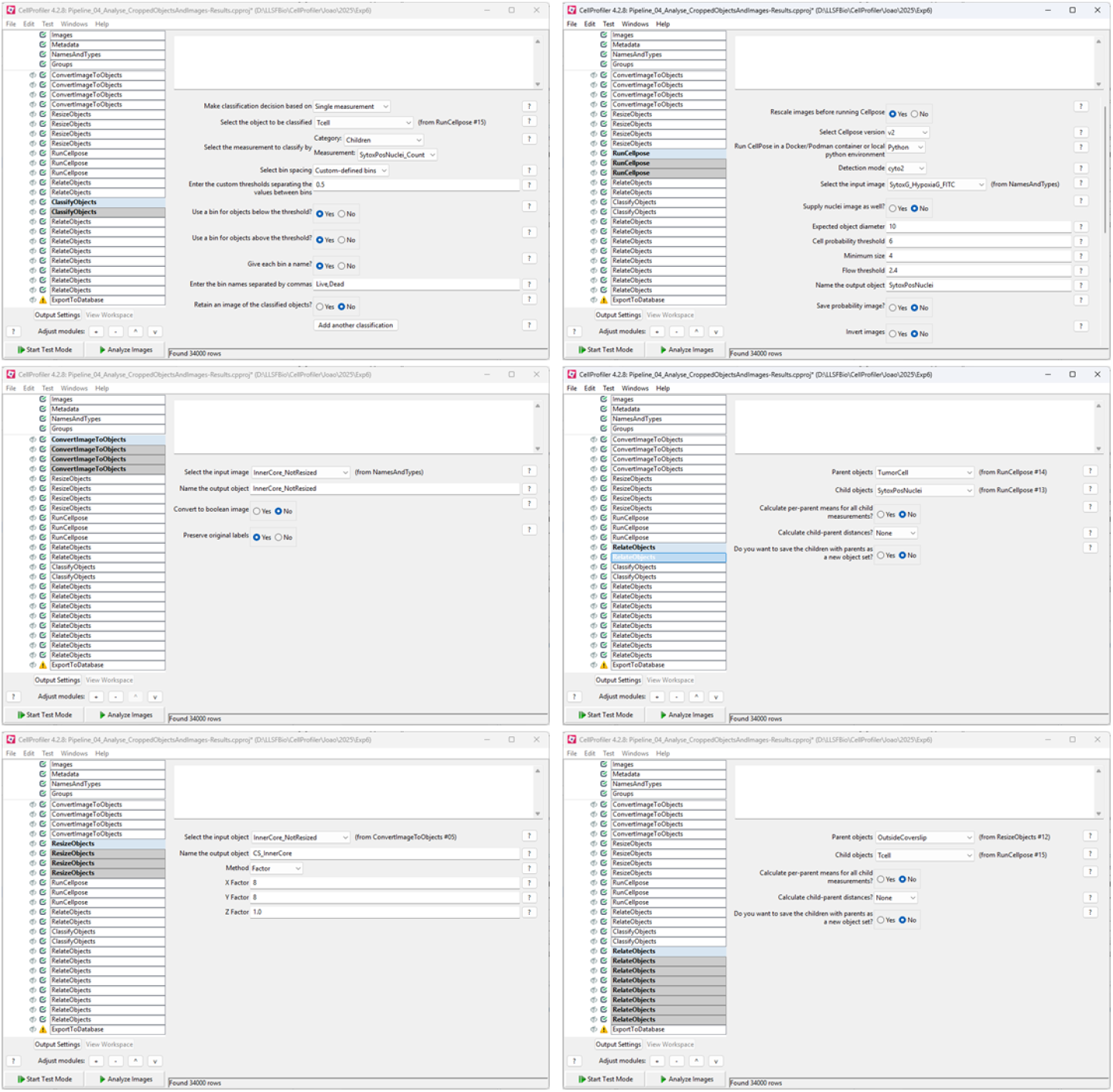
Structure and workflow of the pipeline “Pipeline_04_Analyse_CroppedObjectsAndImages-Results.cpproj”. In **ConvertImageToObjects** and **ResizeObjects**, each cropped ROI is converted into a measurable object and expanded to match its original image scale. Subsequently, **RunCellPose** modules perform AI-GPU-powered segmentation of Raji tumor cells (Cy5), CAR-T cells (CellTrace Violet), and Sytox-positive nuclei of dead cells. **RelateObjects** modules associate Sytox-positive nuclei with corresponding Raji or CAR-T cells and **ClassifyObjects** assigns “live” or “dead” labels based on the identified relation with Sytox nuclei. The final **RelateObjects** link each cell type (Raji or CAR-T) to its spatial ROI (InnerCore, OuterCore, Periphery, Outside) and **ExportToDatabase** modules integrate all cellular features and spatial metadata, generating a unified dataset for downstream analysis of CAR-T function and tumor cell death across hypoxia gradients.

**Supplementary Script 1.**
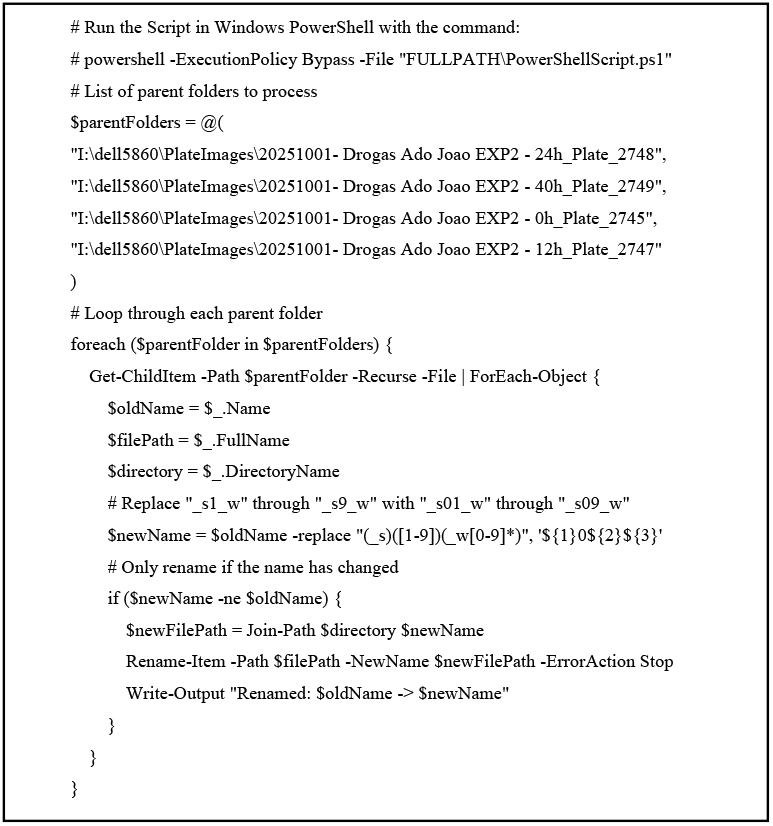
PowerShell script used to automate the renaming of image site numbers across multiple experiment folders for *CellProfiler* batch processing. The script RenameImageSiteNumbers_PowerShellScript_MultipleFolders.ps1 iteratively scans subdirectories containing multi-site image sets, reassigning standardized site identifiers within filenames (e.g., s01–s25) to ensure compatibility with *CellProfiler* metadata extraction and grouping. This automation streamlines preprocessing in high-content screening (HCS) experiments, enabling uniform data organization and reproducible image analysis workflows.

## Bibliography

1. Burugu et al. Semin Cancer Biol (2018). 10.1016/j.semcancer.2017.10.001

2. Doi et al. Oncol Rep (2017). 10.3892/or.2017.5399

3. Azad et al. EMBO Mol Med (2017). 10.15252/emmm.201606674

4. Wang et al. Signal Transduct Target Ther (2020). 10.1038/s41392-020-0144-8

5. Wei et al. Cancer Discov (2018). 10.1158/2159-8290.CD-18-0367

6. Hoffner et al. J Adv Pract Oncol (2019). 10.6004/jadpro.2019.10.4.5

7. Gomes de Morais et al. Curr Oncol Rep (2022). 10.1007/s11912-022-01218-y

8. Shah et al. Signal Transduct Target Ther (2021). 10.1038/s41392-021-00823-w

9. Mao et al. Expert Rev Mol Med (2022). 10.1017/erm.2021.32

10. Titov et al. Cancers (Basel) (2021). 10.3390/cancers13040743

11. Feins et al. Am J Hematol (2019). 10.1002/ajh.25418

12. Salzer et al. Nat Commun (2020). 10.1038/s41467-020-17970-3

13. Tian et al. Journal of hematology & oncology (2020). 10.1186/s13045-020-00890-6

14. Tokarew et al. Br J Cancer (2019). 10.1038/s41416-018-0325-1

15. Zhao et al. Front Immunol (2019). 10.3389/fimmu.2019.02250

16. Bair et al. Am J Hematol (2019). 10.1002/ajh.25457

17. Frey. Am J Hematol (2019). 10.1002/ajh.25442

18. Levin et al. Am J Hematol (2019). 10.1002/ajh.25403

19. Bruni et al. Nat Rev Cancer (2020). 10.1038/s41568-020-0285-7

20. Sitkovsky. Current opinion in pharmacology (2020). 10.1016/j.coph.2020.07.011

21. Sitkovsky. Cancer Discov (2020). 10.1158/2159-8290.CD-19-1280

22. Pai et al. Cancer Treat Res (2020). 10.1007/978-3-030-38862-1_6

23. Hegde et al. Clin Cancer Res (2016). 10.1158/1078-0432.CCR-15-1507

24. Chen et al. Nature (2017). 10.1038/nature21349

25. Joyce et al. Science (2015). 10.1126/science.aaa6204

26. Chavez et al. Ther Adv Hematol (2019). 10.1177/2040620719841581

27. Rossi et al. Cancer Research (2019). 10.1158/1538-7445.Am2019-Ct153

28. Rossi et al. Cancer Research (2018). 10.1158/1538-7445.Am2018-Lb-016

29. Galon et al. Journal of Clinical Oncology (2017). 10.1200/JCO.2017.35.15_suppl.3025

30. Yan et al. Clin Cancer Res (2019). 10.1158/1078-0432.CCR-19-0101

31. Chen et al. JCI Insight (2020). 10.1172/jci.insight.134612

32. Andrea et al. Front Immunol (2022). 10.3389/fimmu.2022.830292

33. Sahai et al. Nat Rev Cancer (2020). 10.1038/s41568-019-0238-1

34. Sunami et al. Cancers (Basel) (2021). 10.3390/cancers13040697

35. Poggi et al. Front Immunol (2018). 10.3389/fimmu.2018.00262

36. Pereira et al. Trends Cancer (2019). 10.1016/j.trecan.2019.09.010

37. Gatenby et al. Br J Cancer (2007). 10.1038/sj.bjc.6603922

38. Park et al. Front Cell Dev Biol (2022). 10.3389/fcell.2022.830208

39. Waibl Polania et al. Front Immunol (2021). 10.3389/fimmu.2021.777073

40. Gowrishankar et al. Mamm Genome (2018). 10.1007/s00335-018-9756-5

41. Rodriguez-Garcia et al. Front Immunol (2020). 10.3389/fimmu.2020.01109

42. Molon et al. Front Immunol (2016). 10.3389/fimmu.2016.00020

43. Daniel et al. Clin Transl Med (2019). 10.1186/s40169-019-0226-9

44. Bartrons et al. J Bioenerg Biomembr (2007). 10.1007/s10863-007-9080-3

45. Hatfield et al. Adv Exp Med Biol (2019). 10.1007/978-3-030-12734-3_8

46. Ben-Shoshan et al. Arterioscler Thromb Vasc Biol (2009). 10.1161/ATVBAHA.108.183319

47. Freemerman et al. J Biol Chem (2014). 10.1074/jbc.M113.522037

48. Hu et al. Front Immunol (2021). 10.3389/fimmu.2021.633361

49. Neo et al. J Clin Invest (2020). 10.1172/JCI128895

50. Anastasiou. Br J Cancer (2017). 10.1038/bjc.2016.412

51. Hobson-Gutierrez et al. Dis Model Mech (2018). 10.1242/dmm.034462

52. Ben-Shoshan et al. Eur J Immunol (2008). 10.1002/eji.200838318

53. Angelin et al. Cell Metab (2017). 10.1016/j.cmet.2016.12.018

54. Schito et al. Trends Cancer (2016). 10.1016/j.trecan.2016.10.016

55. Strickaert et al. Oncogene (2017). 10.1038/onc.2016.411

56. Vander Heiden et al. Science (2009). 10.1126/science.1160809

57. Khalaf et al. Front Immunol (2021). 10.3389/fimmu.2021.656364

58. Ni et al. Front Cell Dev Biol (2021). 10.3389/fcell.2021.637675

59. Bussard et al. Breast Cancer Res (2016). 10.1186/s13058-016-0740-2

60. Labani-Motlagh et al. Front Immunol (2020). 10.3389/fimmu.2020.00940

61. Soongsathitanon et al. J Immunol Res (2021). 10.1155/2021/8840066

62. Hill et al. Semin Cancer Biol (2020). 10.1016/j.semcancer.2019.07.028

63. Zlotnik. Int J Cancer (2006). 10.1002/ijc.2202464.

64. Muller et al. Nature (2001). 10.1038/35065016

65. Boulais et al. Blood (2015). 10.1182/blood-2014-09-570192

66. Kandarakov et al. Int J Mol Sci (2022). 10.3390/ijms23084462

67. Morrison et al. Nature (2014). 10.1038/nature12984

68. Muz et al. Mol Cancer Res (2014). 10.1158/1541-7786.MCR-14-0028

69. Chen et al. J Cell Mol Med (2021). 10.1111/jcmm.17027

70. He et al. Oncol Lett (2021). 10.3892/ol.2021.12953

71. Jiang et al. Transl Pediatr (2021). 10.21037/tp-21-86

72. Bruno et al. Int J Mol Sci (2021). 10.3390/ijms22136857

73. Eisenberg et al. Cancer Lett (2020). 10.1016/j.canlet.2020.04.016

74. Gorlach. Circ Res (2005). 10.1161/01.RES.0000174112.36064.77

75. Lim To et al. Placenta (2015). 10.1016/j.placenta.2015.04.005

76. Antonioli et al. Trends Mol Med (2013). 10.1016/j.molmed.2013.03.005

77. Eltzschig et al. J Exp Med (2003). 10.1084/jem.20030891

78. Linden et al. Annual review of immunology (2019). 10.1146/annurev-immunol-051116-052406

79. Baldwin et al. Pflugers Arch (2004). 10.1007/s00424-003-1103-2

80. Casanello et al. Circ Res (2005). 10.1161/01.RES.0000172568.49367.f8

81. Boison et al. Cancer Cell (2019). 10.1016/j.ccell.2019.10.007

82. Dong et al. J Immunol (1996). https://www.ncbi.nlm.nih.gov/pubmed/8568233

83. Ohta et al. Proc Natl Acad Sci U S A (2006). 10.1073/pnas.0605251103

84. Zarek et al. Blood (2008). 10.1182/blood-2007-03-081646

85. Stagg et al. Proc Natl Acad Sci U S A (2010). 10.1073/pnas.0908801107

86. Beavis et al. Proc Natl Acad Sci U S A (2013). 10.1073/pnas.1308209110

87. Desmet et al. Proc Natl Acad Sci U S A (2013). 10.1073/pnas.1222085110

88. Sitkovsky. Trends Immunol (2009). 10.1016/j.it.2008.12.002

89. Allard et al. Nat Rev Clin Oncol (2020). 10.1038/s41571-020-0382-2

90. Leone et al. J Immunother Cancer (2018). 10.1186/s40425-018-0360-8

91. Franco et al. Cells (2021). 10.3390/cells10112831

92. Moesta et al. Nat Rev Immunol (2020). 10.1038/s41577-020-0376-4

93. Kotulova et al. Int J Mol Sci (2021). 10.3390/ijms222212569

94. Augustin et al. J Immunother Cancer (2022). 10.1136/jitc-2021-004089

95. Vickman et al. Oncotarget (2020). 10.18632/oncotarget.27736

96. Ringquist et al. Adv Drug Deliv Rev (2021). 10.1016/j.addr.2021.114003

97. Hammel et al. Adv Drug Deliv Rev (2022). 10.1016/j.addr.2022.114111

98. Ando et al. Acta Biomater (2021). 10.1016/j.actbio.2021.03.076

99. Pietrobon et al. J Transl Med (2021). 10.1186/s12967-020-02667-4

100. Pietrobon et al. Front Immunol (2020). 10.3389/fimmu.2020.604967

101. Carmona-Fontaine et al. Proc Natl Acad Sci U S A (2017). 10.1073/pnas.1700600114

102. Janska et al. Dis Model Mech (2021). 10.1242/dmm.048942

103. Carmona-Fontaine et al. Proc Natl Acad Sci U S A (2013). 10.1073/pnas.1311939110

104. Ando et al. Adv Healthc Mater (2019). 10.1002/adhm.201900001

105. Spoerri et al. Front Digit Health (2021). 10.3389/fdgth.2021.668390

106. Zboralski et al. Methods Mol Biol (2019). 10.1007/978-1-4939-8885-3_19

107. Hoellenriegel et al. Blood (2014). 10.1182/blood-2013-03-493924

108. Steurer et al. Haematologica (2019). 10.3324/haematol.2018.205930

109. Zboralski et al. Cancer Immunol Res (2017). 10.1158/2326-6066.CIR-16-0303

110. Zboralski et al. Blood (2016). 10.1182/blood.V128.22.3021.3021

111. Jiang et al. Small (2021). 10.1002/smll.202004282

112. Pulvertaft. Lancet (1964). 10.1016/s0140-6736(64)92345-1

113. Takakuwa et al. Am J Pathol (2004). 10.1016/S0002-9440(10)63184-7

114. Kamio et al. Blood (2003). 10.1182/blood-2002-12-3656

115. Schwenk et al. Blut (1975). 10.1007/BF01634146

116. Gillis et al. J Exp Med (1980). 10.1084/jem.152.6.1709

117. Willingham et al. Cancer Immunol Res (2018). 10.1158/2326-6066.CIR-18-0056

118. Fong et al. Cancer Discov (2020). 10.1158/2159-8290.CD-19-0980

119. Brunner et al. Immunology (1968). https://www.ncbi.nlm.nih.gov/pubmed/4966657 https://www.ncbi.nlm.nih.gov/pmc/articles/PMC1409286/pdf/immunology00397-0036.pdf

120. Decker et al. J Immunol Methods (1988). 10.1016/0022-1759(88)90310-9

121. Neri et al. Clin Diagn Lab Immunol (2001). 10.1128/CDLI.8.6.1131-1135.2001

122. Karimi et al. PloS one (2014). 10.1371/journal.pone.0089357

123. Matta et al. Sci Rep (2018). 10.1038/s41598-017-18606-1

124. Rossignol et al. MAbs (2017). 10.1080/19420862.2017.1286435

125. Cerignoli et al. PloS one (2018). 10.1371/journal.pone.0193498

126. Martinez-Serra et al. Onco Targets Ther (2014). 10.2147/OTT.S62887

127. Jedema et al. Blood (2004). 10.1182/blood-2003-06-2070

128. Hermans et al. J Immunol Methods (2004). 10.1016/j.jim.2003.10.017

129. Maldini et al. J Immunol Methods (2020). 10.1016/j.jim.2020.112830

130. Sant et al. Drug Discov Today Technol (2017). 10.1016/j.ddtec.2017.03.002

131. Colombo et al. Int J Mol Sci (2021). 10.3390/ijms22041633

132. Rodrigues et al. Trends Cancer (2021). 10.1016/j.trecan.2020.10.009

133. Franchi-Mendes et al. Cancers (Basel) (2021). 10.3390/cancers13184610

134. Gonzales et al. N Biotechnol (2009). 10.1016/j.nbt.2009.06.982

135. Fouda et al. J Vis Exp (2017). 10.3791/55199

136. Jutz et al. J Immunol Methods (2016). 10.1016/j.jim.2016.01.007

137. Jutz et al. Oncotarget (2017). 10.18632/oncotarget.17615

138. Salik et al. Journal of hematology & oncology (2020). 10.1186/s13045-020-00947-6

139. Rydzek et al. Mol Ther (2019). 10.1016/j.ymthe.2018.11.015

140. Rosskopf et al. Oncotarget (2018). 10.18632/oncotarget.24807

141. Muller et al. Clin Transl Immunology (2020). 10.1002/cti2.1216

142. Morimoto et al. Oncotarget (2018). 10.18632/oncotarget.26139

143. Fierle et al. Sci Rep (2022). 10.1038/s41598-022-05058-5

